# Multi-way modelling of oral microbial dynamics and host-microbiome interactions during induced gingivitis

**DOI:** 10.1101/2024.03.18.585469

**Authors:** G.R. van der Ploeg, B.W. Brandt, B.J.F. Keijser, M.H. van der Veen, C.M.C. Volgenant, E. Zaura, A.K. Smilde, J.A. Westerhuis, A. Heintz-Buschart

**Author notes:** Author for correspondence: Anna Heintz-Buschart.

## Abstract

Gingivitis - the inflammation of the gums - is a reversible stage of periodontal disease. It is caused by dental plaque formation due to poor oral hygiene. However, gingivitis susceptibility involves a complex set of interactions between the oral microbiome, oral metabolome and the host. In this study, we investigated the dynamics of the oral microbiome and its interactions with the salivary metabolome during experimental gingivitis in a cohort of 41 systemically healthy participants. We use Parallel Factor Analysis (PARAFAC), which is a multi-way generalization of Principal Component Analysis (PCA) that can model the variability in the response due to subjects, variables and time. Using the modelled responses, we identified microbial subcommunities with similar dynamics that connect to the magnitude of the gingivitis response. By performing high level integration of the predicted metabolic functions of the microbiome and salivary metabolome, we identified pathways of interest that describe the changing proportions of Gram-positive and Gram-negative microbiota, variation in anaerobic bacteria, biofilm formation and virulence.

## Introduction

Microbial oral diseases, such as dental caries and gum inflammation, are a global public health burden (Peres et al., 2019; Petersen et al., 2005; WHO, 2022). Together, they are the most prevalent health condition world-wide, impacting quality of life and imposing a significant economic burden with 5-10% of public health expenditure in most industrialized countries (WHO, 2022; Widström et al., 2004). Prevention of oral diseases is therefore of critical importance.

Gingivitis is a reversible stage of periodontal disease marked by inflamed, bleeding gums (Chapple et al., 2015; Curtis et al., 2020; Huang et al., 2014). It is caused by dental plaque accumulation due to poor oral hygiene (Löe et al., 1965; Murakami et al., 2018) and is present in a substantial part of the global population (Han et al., 2020; WHO, 2022). The pathogenesis of gingivitis involves a complex set of interactions between the oral microbiome, the oral metabolome and the host (Chapple et al., 2018; Huang et al., 2014). These are in turn dependent on many factors (Chapple et al., 2018; Kilian et al., 2016), including host diet (Van Der Velden et al., 2011), life-style (Bergström & Preber, 1986), and genetics (Nibali et al., 2017). Therefore, the oral microbiome alone does not determine whether and to what extent gingivitis develops, as can be seen by the high degree of individual variation in gingivitis susceptibility seen in experimental gingivitis models after refraining from toothbrushing (Axelsson et al., 1991; Chapple et al., 2015, 2018; Guk et al., 2020; Huang et al., 2014; van der Veen et al., 2016)

While there is no monocausal microbial agent in gingivitis, the gingival crevice microbiota underneath the gums shifts to a dysbiotic state (Abusleme et al., 2021; Curtis et al., 2020; Diaz et al., 2016). The depletion of Gram-positive bacteria, such as *Rothia dentocariosa,* and enrichment in Gram-negative bacteria such as *Porphyromonas gingivalis*, *Prevotella* spp. and *Selenomonas* spp. is frequently observed in case-control comparisons, despite this shift being less pronounced compared to periodontitis progression (Curtis et al., 2020; Huang et al., 2014; Kistler et al., 2013; Schincaglia et al., 2017). Additionally, increasing subgingival anaerobiosis during gingivitis progression promotes the growth of pathobionts (Curtis et al., 2020; Diaz et al., 2002; Ter Steeg et al., 1988).

Being the medium through which the host and the microbiota interact, the salivary metabolome is often studied as a non-invasive diagnostic of oral disease (Dawes & Wong, 2019; Proctor, 2016). A well-known host-microbiome interaction is the relation between host sugar consumption and the production of acids by the dental plaque microbiota. This lowers the local pH and shifts the microbiota further towards dysbiosis (König & Navia, 1995; Marsh, 1991). However, many other dynamic microbe-microbe and host-microbiome interactions that occur throughout gingivitis onset and progression are not well understood.

Here, we investigate the spatially resolved oral bacterial microbiome and its interactions with the salivary metabolome during experimental gingivitis to (1) identify microbial subcommunities with common dynamics and (2) pinpoint potential impacts on biochemical pathways. Reversible gingivitis was induced in a cohort of 41 systemically healthy participants by omission of oral hygiene during a two-week gingivitis challenge (**Figure 1A**; (Prodan et al., 2016; van der Veen et al., 2016)). Previously reported plaque levels, gingival bleeding upon probing, and red fluorescent plaque quantification allowed grouping of the participants into identified low, mid and high responders based on the last day of the intervention (**Figure 1B**; (van der Veen et al., 2016)). During a two-week baseline period at two time points (day −14 and day 0), a challenge period at four time points (day 2 to day 14), and one week resolution period at one time point (day 21), the oral microbiome was determined at six sites: tongue, saliva, supragingival plaque at the lower and upper lingual surfaces and supragingival plaque at the lower and upper interproximal surfaces (**Figure 1C**). To identify the individual variation in the time-resolved response of the oral microbiome to the challenge and describe commonly responding (groups of) microbiota, we employ Parallel Factor Analysis (PARAFAC; (Carroll & Chang, 1970; Harshman, 1970)), for modelling the subjects, time, and microbial abundances at each site (**Figure 1D**). The salivary metabolome dynamics, assessed in unstimulated saliva during the challenge, is likewise modelled and integrated with microbiome models at the biochemical pathway level. These analyses should yield insights into microbial processes that underlie individual gingivitis responses.

**Figure 1:**
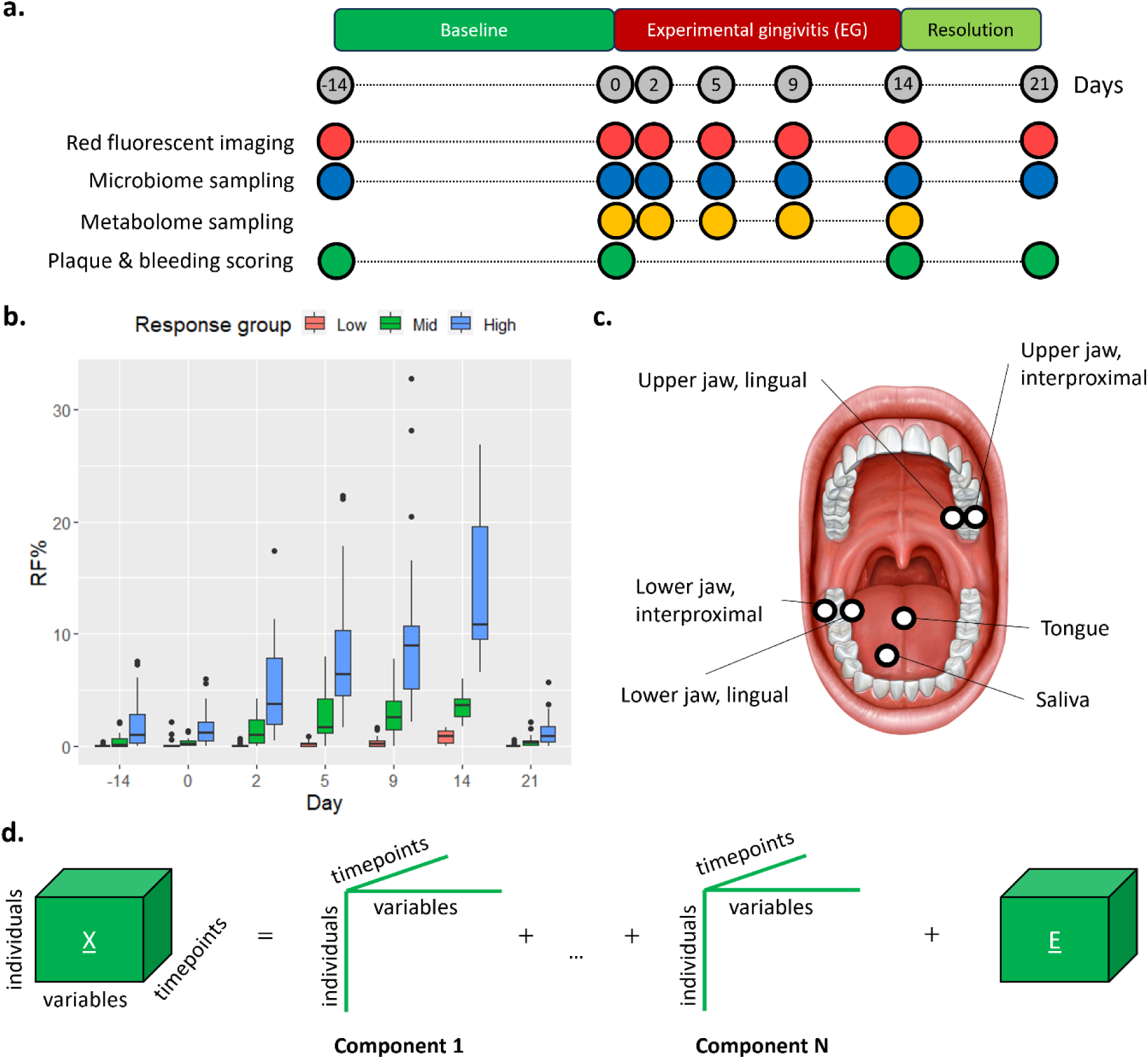
Overview of the study design and analysis. (a) Timeline of experimental gingivitis challenge with analyses; (b) gingivitis responses: Three groups of responders were identified based on red fluorescent dental plaque (RF%) at day 14: low (0-1.7%), mid (1.8-6.0%) and high (6.0%+); (c) oral sampling sites; (d) data analysis approach: Parallel factor analysis (PARAFAC), a multi-way generalization of Principal Component analysis, decomposes a multi-way data cube into components with three loading vectors each: one for the subjects, one for the variables (here: microbial relative abundances) and one for time. This approach allows us to identify the individual variation in the time-resolved response to the challenge and to describe commonly responding groups of microbiota.

## Results

### PARAFAC describes clinically relevant variation in oral microbiomes between subjects

To describe the microbial dynamics of each oral site for all subjects across time, we analysed 16S rRNA gene amplicon sequencing data for a total of 14,014 amplicon sequence variants (ASVs). After removal of sparse ASVs, we created separate PARAFAC models for each oral site, choosing the most appropriate number of PARAFAC components by inspecting the number of iterations needed to converge (Bro, 1997), the core consistency diagnostic (Bro & Kiers, 2003), the variation explained (Bro, 1997), and the tucker congruence coefficient per mode (Lorenzo-Seva & ten Berge, 2006; Tucker, 1951) (**Supplementary Data**). This yielded two-component models for the upper jaw lingual plaque (19.7% explained variation) and upper jaw interproximal plaque (16.5%), tongue (31.5%) and saliva (17.5%) microbiomes (**Figure 2**). Lower jaw lingual and interproximal plaque microbiomes were best represented by one-component models (explaining 11.4% and 7.2% of the variation, respectively). The model of the lower jaw interproximal plaque microbiome was not investigated further due to the limited amount of explained variation and due to the model not describing biologically meaningful information.

**Figure 2:**
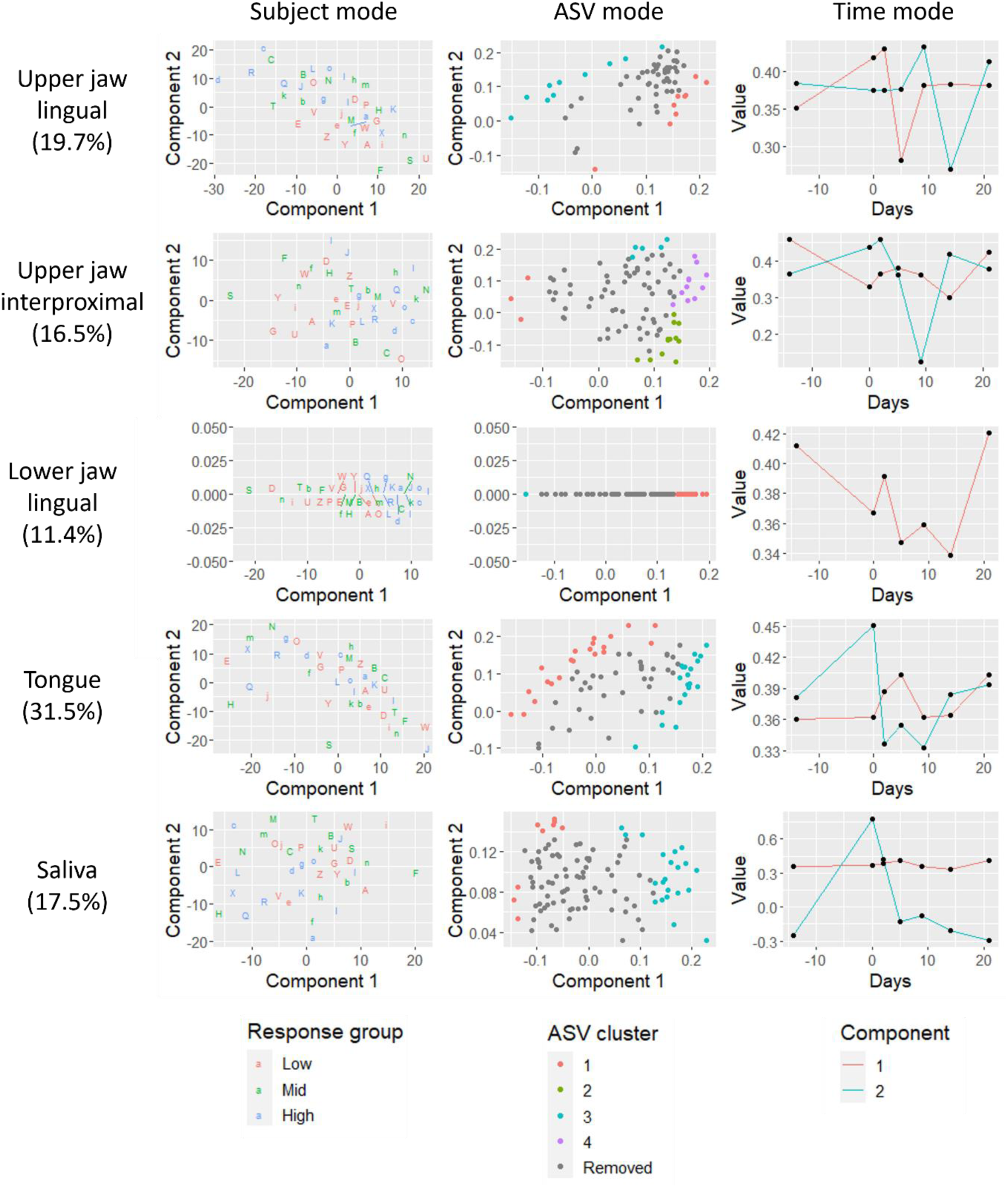
Overview of the PARAFAC models per sample type. In the rows, the PARAFAC models per sample type are described: upper jaw lingual plaque, upper jaw interproximal plaque, lower jaw lingual plaque, tongue and saliva. Variance explained per model are shown in the sample type labels. In the left column the loadings of the first and second component in the subject mode are plotted against each other, with every subject having a unique character to identify them across the sample types. In the middle column the loadings of the first and second component in the ASV mode are plotted against each other. We identified microbial subcommunities (ASV clusters) with common responses per sampling site by selecting and then clustering well-modelled ASVs based on their fitted response (Methods, **Table 2**). ASVs are color-coded by cluster number (or shown in grey if removed prior to clustering). In the right column, the loadings per time point are shown per component. The model corresponding to the lower jaw lingual plaque only has one component. The model of the lower jaw interproximal plaque microbiome was not investigated further due to the limited amount of explained variation and due to the model not describing biologically meaningful information.

To interpret the models, we tested the correlation between the time-resolved subject loadings of each component with the measured parameters of gingivitis. This approach revealed correlation of components with one or more clinical parameter for the three plaque microbiomes (**Table 1**). Overall, the PARAFAC models describe the variation that exists between subjects, rather than the variation within a subject (one-sided Wilcoxon rank-sum test: p=0.020, **Supplementary Figures A1 and A2**). Furthermore, subject age was found to be significantly described by the first component of the model corresponding to the tongue microbiome samples (p=0.035) and the second component of the model corresponding to the saliva microbiome samples (p=0.038) (**Supplementary Table B1**). Gender was not significantly described by any model (**Supplementary Table B1**). We also performed Pearson correlation tests of the time-resolved subject loadings per component with richness and evenness. All microbiome models were found to contain at least one component that significantly described richness or evenness (p≤0.05; **Table 1**). In conclusion, the PARAFAC models represented clinically and ecologically relevant parameters.

**Table 1:**
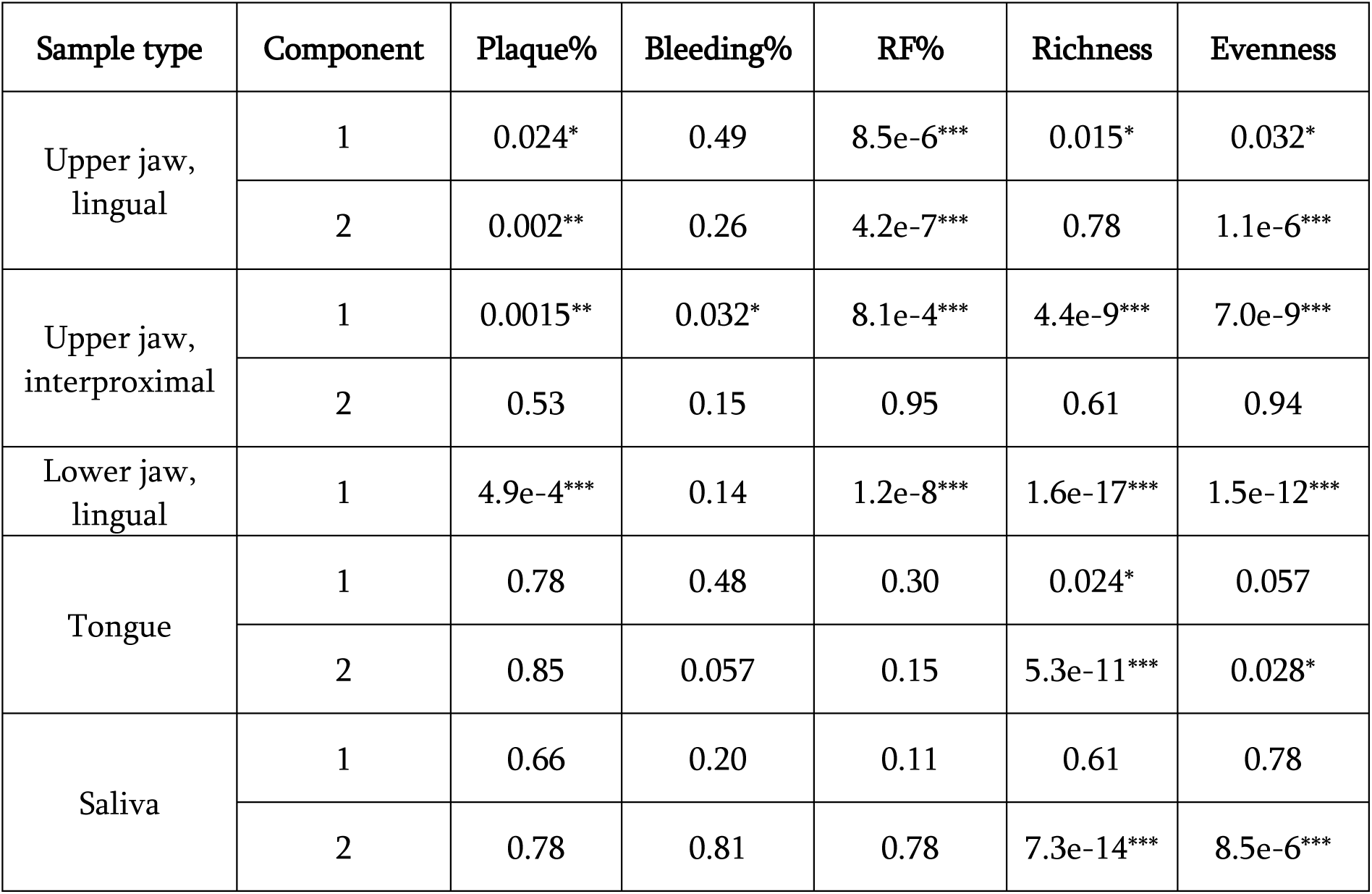
Overview of the correlation test results between the time-resolved subject loadings of the PARAFAC models and the clinical parameters of gingivitis and microbiome diversity metrics. Red fluorescence was assessed at every time point. Plaque and bleeding scores were only assessed at day −14, 0, 14 and 21 of the study. Plaque%: the percentage of sites in the mouth that were covered in plaque. Bleeding%: the percentage of sites in the mouth that bled upon probing. RF%: the percentage of sites that fluoresced red. Richness: the number of nonzero counts in a sample. Evenness: Shannon diversity divided by log(richness). Benjamini-Hochberg corrected p-values of Pearson correlation tests between the subject loadings of one component and the clinical parameters are reported (*: p≤0.05; **: p≤0.01; ***: p≤0.001).

| Sample type | Component | Plaque% | Bleeding% | RF% | Richness | Evenness |
| --- | --- | --- | --- | --- | --- | --- |
| Upper jaw,<br>lingual | 1 | 0.024* | 0.49 | 8.5e-6*** | 0.015* | 0.032* |
|  | 2 | 0.002** | 0.26 | 4.2e-7*** | 0.78 | 1.1e-6*** |
| Upper jaw,<br>interproximal | 1 | 0.0015** | 0.032* | 8.1e-4*** | 4.4e-9*** | 7.0e-9*** |
|  | 2 | 0.53 | 0.15 | 0.95 | 0.61 | 0.94 |
| Lower jaw,<br>lingual | 1 | 4.9e-4*** | 0.14 | 1.2e-8*** | 1.6e-17*** | 1.5e-12*** |
| Tongue | 1 | 0.78 | 0.48 | 0.30 | 0.024* | 0.057 |
|  | 2 | 0.85 | 0.057 | 0.15 | 5.3e-11*** | 0.028* |
| Saliva | 1 | 0.66 | 0.20 | 0.11 | 0.61 | 0.78 |
|  | 2 | 0.78 | 0.81 | 0.78 | 7.3e-14*** | 8.5e-6*** |

### PARAFAC describes microbial subcommunities with common gingivitis responses

Building on the interpretability of the subject modes (**Table 1**), we identified microbial subcommunities with common responses per sampling site by selecting and then clustering well-modelled ASVs based on their fitted response (**Figure 2**, for cluster membership see **Table 2**). Per sampling site, at least one subcommunity containing mainly pathobionts or pro-inflammatory genera such as *Actinomyces*, *Campylobacter*, *Capnocytophaga*, *Fusobacterium*, *Leptotrichia* and *Porphyromonas* (Huang et al., 2014; Kirst et al., 2015; Kistler et al., 2013; Schincaglia et al., 2017; The Human Microbiome Project Consortium, 2012) and one subcommunity containing mainly commensal genera such as *Kingella oralis* (Abusleme et al., 2021), *Streptococcus* spp. (Caporaso et al., 2011; Zaura et al., 2009), and *Veillonella* spp. (Moore & Moore, 1994) have been identified.

**Table 2:** Overview of all identified species per ASV cluster. ASVs within the same cluster show a similar response over time to the gingivitis intervention. Clustered ASVs are only reported here if two criteria were met: (1) the taxonomic classification was resolved to species level by DADA2 using the SILVA (v138) and HOMD (v15.22) databases and (2) the annotation agreed with a separate best hit annotation run using the HOMD (v15.22) database. Cases where the classification was resolved to species level by the DADA2 run but only to the genus level by the separate HOMD run are also reported.

| Sample type | Cluster | Number of members | Identified species |
| --- | --- | --- | --- |
| Upper jaw, lingual | 1 | 9 | *Actinomyces naeslundii/oris/viscosus*, *Kingella oralis*, *Rothia dentocariosa*, *Streptococcus cristatis/sinensis*, *Veillonella dispar/parvula*, *Veillonella atypica/dispar*, *Veillonella parvula/rogosae/tobetsuensis* |
|  | 2 | 10 | *Fusobacterium canifelinus/nucleatum*, *Fusobacterium massiliense*, *Granulicetella elegans*, *Prevotella nanceiensis*, *Veillonella massiliensis* |
| Upper jaw, interproximal | 1 | 3 | *Rothia dentocariosa*, *Veillonella atypica/dispar* |
|  | 2 | 12 | *Abiotrophia defectiva*, *Aggregatibacter aphrophilus/kilianii*, *Capnocytophaga sputigena*, *Lachnoanaerobaculum cf.*, *Lautropia mirabilis*, *Rothia aeria/dentocariosa* |
|  | 3 | 5 | *Corynebacterium matruchotii*, *Leptotrichia hofstadii*, *Leptotrichia shahii/wadei* |
|  | 4 | 11 | *Aggregatibacter segnis*, *Capnocytophaga granulosa*, *Capnocytophaga leadberri*, *Cardiobacterium hominis*, *Fusobacterium canifelinus/nucleatum*, *Fusobacterium nucleatum*, *Leptotrichia buccalis*, *Porphyromonas pasteri* |
| Lower jaw, lingual | 1 | 21 | *Actinomyces georgiae/pacaensis*, *Aggregatibacter segnis*, *Campylobacter concisus*, *Capnocytophaga granulosa*, *Capnocytophaga leadbetteri*, *Cardiobacterium hominis*, *Corynebacterium matruchotii*, *Fusobacterium canifelinus/nucleatum*, *Fusobacterium nucleatum*, *Fusobacterium hwasooki/nucleatum/periodonticum*, *Lachnoanaerobaculum cf.*, *Leptotrichia hofstadii*, *Porphyromonas pasteri* |
|  | 2 | 1 | *Streptococcus salivarius* (putative) |
| Tongue | 1 | 21 | *Actinomyces lingnae/marseillensis/pacaensis*, *Capnocytophaga leadbetteri*, *Fusobacterium periodonticum*, *Granulicatella adiacens/para-adiacens*, *Lachnoanaerobaculum cf.*, *Oribacterium parvum*, *Oribacterium sinus*, *Peptostreptococcus stomatis*, *Porphyromonas pasteri*, *Prevotella nanceiensis*, *Veillonella parvula/rogosae/tobetsuensis* |
|  | 2 | 20 | *Actinomyces graevenitzi*, *Alloprevotella rava*, *Atopobium parvulum*, *Lachnoanaerobaculum orale/saburreum*, *Prevotella histicola*, *Prevotella jejuni/melaninogenica*, *Prevotella salivae*, *Prevotella veroralis*, *Rothia mucilaginosa*, *Stomatobaculum longum* |
| Saliva | 1 | 9 | *Granulicatella adiacens/para-adiacens*, *Porphyromonas pasteri*, *Rothia aeria/dentocariosa* |
|  | 2 | 21 | *Actinomyces graevenitzi*, *Atopobium parvulum*, *Campylobacter concisus*, *Lachnoanaerobaculum orale/saburreum*, *Prevotella histicola*, *Prevotella jejuni/melanogenica*, *Prevotella salivae*, *Rothia mucilaginosa*, *Stomatobaculum longum*, *Veillonella atypica/dispar* |

The sum of the relative abundances of the ASVs in each cluster was determined to test the difference between the microbiomes of individuals in the low and high response groups at every time point per oral site (Benjamini-Hochberg corrected permutation test of mean difference: p≤0.05; **Supplementary Table B2)**. This approach revealed significant differences in the relative abundance of most plaque-associated ASV clusters in high responders compared to the low responders at baseline (day −14; **Figure 3**, **Supplementary Figure A3**). As such, the PARAFAC models of plaque microbiomes describe microbial subcommunities with common dynamics that connect to the magnitude of the gingivitis response.

**Figure 3:**
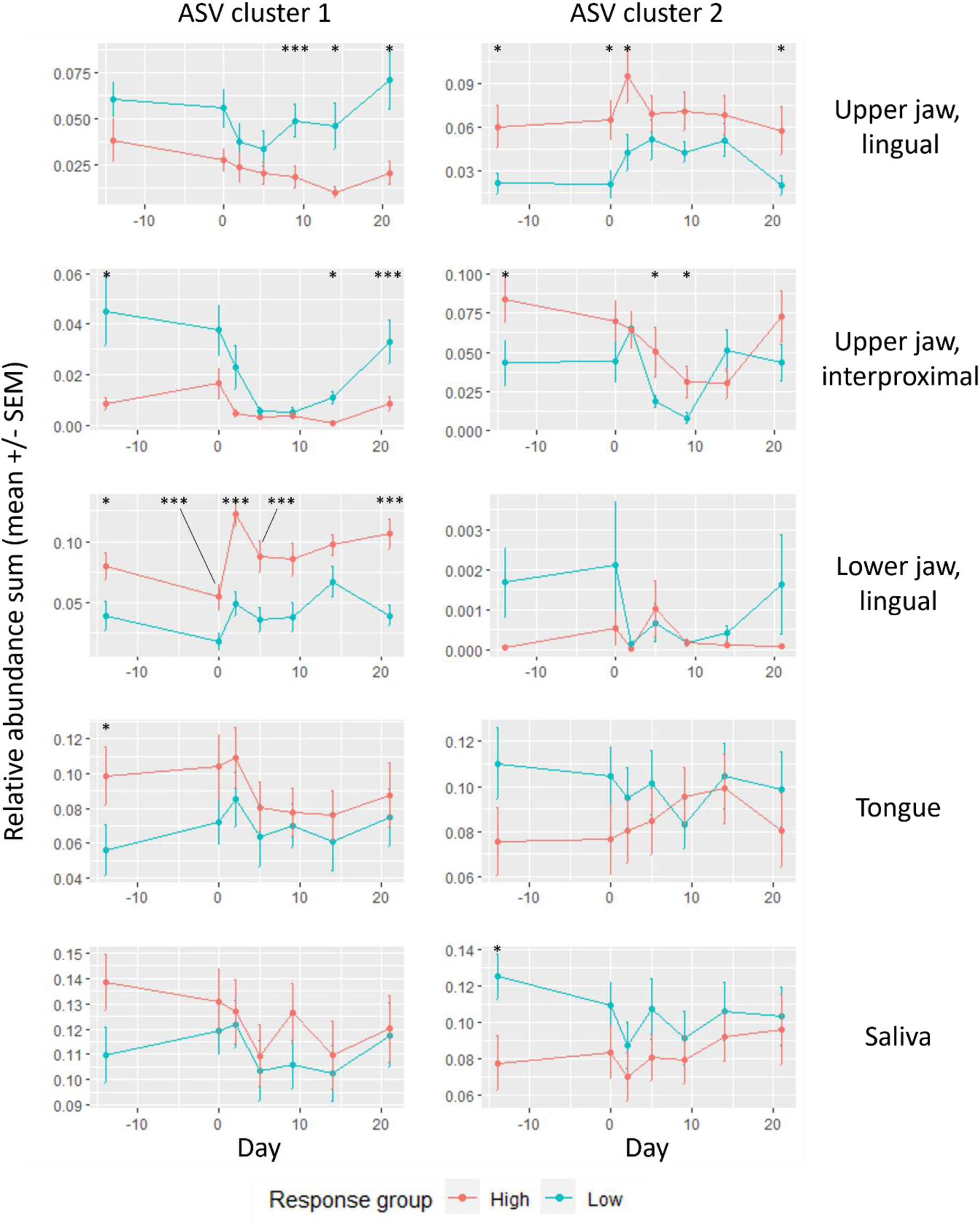
Overview of the sums of relative abundances per ASV cluster and sample type, separating subjects by response group. Error bars correspond to the standard error of the mean (SEM). The mean difference between the high and low response groups per time point was tested using a Benjamini-Hochberg corrected permutation test of 999 iterations (*: p≤0.05; **: p≤0.01; ***: p≤0.001). Mid responders and ASV clusters 3 and 4 of the upper jaw interproximal plaque samples are not shown for visual clarity (**Supplementary Figure A3** and **A4**, respectively). Please refer to **Table 2** for the list of identified species per ASV cluster.

We further investigated the robustness of the ecosystem during gingivitis onset and progression by comparing the sum of the relative abundances per cluster at the start and end of the intervention (**Supplementary Table B3**). We found significant differences between the relative abundances of some ASV clusters at the start and end of the gingivitis challenge in lower jaw lingual (ASV cluster 1) and upper jaw interproximal (ASV cluster 4) plaque microbiomes in all response groups (Benjamini-Hochberg corrected two-sided Wilcoxon rank sum test: p≤7.2e-13 and p≤2.6e-6, respectively). For a further three ASV clusters (upper jaw lingual ASV cluster 1 and upper jaw interproximal clusters 1 and 3), we found the sum of relative abundances to be significantly different (p=0.002, p=0.0025, p=0.0039, respectively) in high responders, but not in low responders. These results suggest that ecosystem stability is different between high and low responders in several oral niches.

### Integration of the microbiome and metabolome PARAFAC models identifies host-microbiome interactions at the pathway level

With the established connection between the clinical parameters of gingivitis and the variation in the plaque microbiomes as modelled by PARAFAC, we investigated host-microbiome interactions at the biochemical pathway level. We obtained functional predictions of the microbiome per oral site using Tax4Fun2 (Aßhauer et al., 2015; Wemheuer et al., 2020). We then created separate PARAFAC models of the salivary metabolomics data and of the functional prediction of the microbiomes at each oral site. Despite correcting for correlation between most salivary metabolite levels, likely due to variable water content of the saliva (see Methods), the subject loadings of both components of the model corresponding to the salivary metabolome were found to correlate with the correcting factor (p=2.0e-14 and p=2.0e-5, respectively). Regardless, we expected the salivary metabolomics model to have sufficient freedom to also describe variation related to the gingivitis response.

The well-modelled molecular functions (KEGG orthologous groups belonging to pathways that produce or consume the salivary metabolites) and well-modelled salivary metabolites were combined in a ranked list. KEGG-pathway enrichment was identified using SetRank, which is a functional enrichment algorithm that corrects for multiple pathway membership (Simillion et al., 2017). This revealed that the microbiome and metabolome responses to the gingivitis challenge reflected differences in carbon and energy metabolism, as well as cell wall structure (**Figure 4, Supplementary Figure A5, Supplementary Table B4**). For example, the tongue microbiome and salivary metabolome were involved in a joint response in the biosynthesis of cofactors and carbon metabolism (p=0.0045 and p=0.0052, respectively). The quorum sensing pathway was significantly enriched in all plaque microbiomes (p≤0.05). The pathway enrichment results therefore reflect the microbiome responses and highlight potential metabolic interactions with the human host.

**Figure 4:**
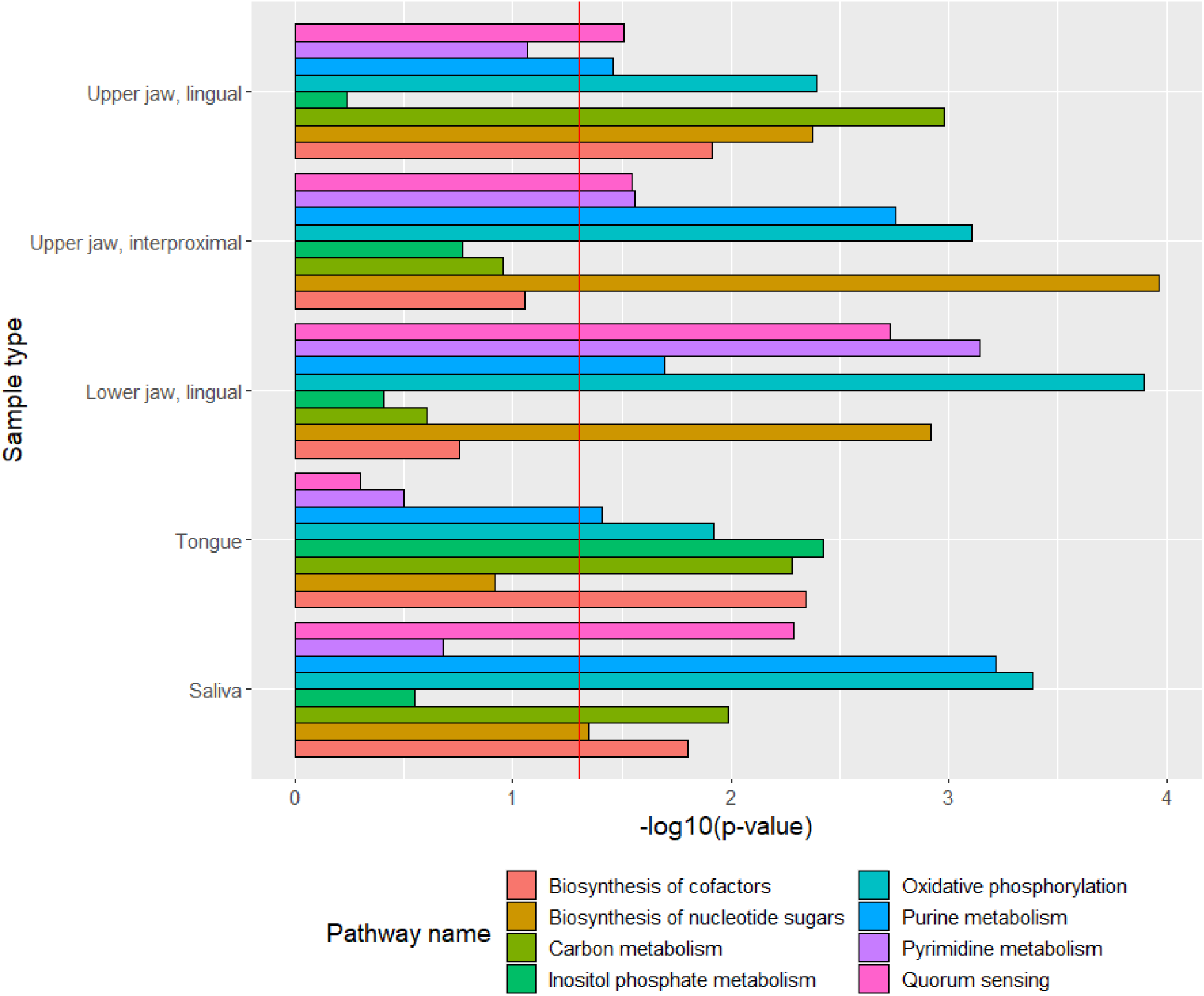
Overview of the pathway enrichment results per combination of oral site microbiome and the salivary metabolome. Tax4Fun2 was used to create functional predictions from ASV data. Well-modelled metabolites and microbiome molecular functions were integrated by mapping them to KEGG pathway level. SetRank was then used to test for pathway enrichment and corrected for multiple-pathway membership. Pathways were filtered to be significant (p<=0.01) in at least one sample type and to contain at least two well-modelled KEGG molecular functions and two well-modelled metabolites. The red vertical line corresponds to p=0.05.

## Discussion

We have shown that the unsupervised modelling approach of PARAFAC mainly described variation between subjects, as expected from an algorithm describing the maximum amount of variation that is present in the data. The modelled variation could be attributed to clinically relevant parameters. In addition, we have observed that the model corresponding to the salivary metabolomics samples partly describes the salivary dilution despite correcting for it using Probabilistic Quotient Normalization (Dieterle et al., 2006). This might be an artifact due to an incorrect assumption that correcting for water content of saliva can be done in the same way as for urine metabolomics samples. Future work on this topic should focus on the appropriateness of using Probabilistic Quotient Normalization prior to performing a decomposition analysis. Other multi-way approaches exist to describe a specific type of variation (Bro, 1997; Carroll & Chang, 1970; Harshman, 1970). For example, in N-way partial least squares (NPLS) the model is constrained to regress on an outcome variable (Bro, 1996). Additionally, the PARAFAC models of the microbiome and metabolome could be created in a linked fashion by keeping one of the modes equal to each other, known as coupled matrix and tensor factorization (Acar et al., 2014, 2015; Singh & Gordon, 2008). Each of these approaches might yield new insights as the models explicitly look at variation that is common between the datasets. Further research is needed to compare such methods to PARAFAC in the context of multi-omics data.

We observed that the microbiome PARAFAC models partly describe richness and evenness. This result gives further evidence of a relationship between oral health and microbial diversity which has been described before (Anderson et al., 2018; Schoilew et al., 2019; Thomas et al., 2014). While the methodology used in this study targeted only bacteria, while other members of the oral microbiome – such as viruses and fungi – might also be relevant in gingivitis onset and progression (Baker et al., 2017; Nobbs & Jenkinson, 2015). We identified several bacterial subcommunities within the oral sites that responded similarly to the intervention (**Table 2**). For example, a group of ASVs was significantly more abundant in individuals in the high responder group compared to low responders and represented species of known pathogenic or pro-inflammatory genera such as *Actinomyces*, *Campylobacter*, *Capnocytophaga*, *Fusobacterium*, *Leptotrichia*, and *Porphyromonas* (Huang et al., 2014; Kirst et al., 2015; Kistler et al., 2013; Schincaglia et al., 2017; The Human Microbiome Project Consortium, 2012). Clusters of ASVs that were significantly more abundant in low responders compared to high responders contained known commensal species related to oral health, such as *Kingella oralis* (Abusleme et al., 2021), *Streptococcus* spp. (Caporaso et al., 2011; Zaura et al., 2009), and *Veillonella* spp. (Moore & Moore, 1994). It has to be noted that sequencing based only on the V4 region of the 16S rRNA gene may not provide sufficient resolution to correctly assign pathogenicity or pro-inflammatory traits all ASVs. Similarly, some ASV have been assigned to bacterial taxa that are not commonly observed in the oral cavity; this may indicate that these bacteria were present in the sample due problematic hygiene in the host, or be due to classification errors. Finally, we observed a difference in ecosystem stability between high and low responders in some parts of the oral cavity. This suggests that in some subjects, the ecosystem can withstand the ecological pressure of plaque accumulation and has some mechanisms that prevents healthy microbiota from decreasing in abundance. Further research is needed to identify these mechanisms and the microbiota involved.

By integrating the PARAFAC models of the salivary metabolomics data and of the predicted metabolic functions from the microbiome data, we identified enriched pathways that reflected the changing proportions of Gram-positive and -negative microbiota (cell membrane/wall component pathways, such as lipopolysaccharide and peptidoglycan biosynthesis). Quorum sensing systems were found to be significantly enriched in all plaque sites likely due to the presence of various *Streptococcus* spp., that carry genes for oligopeptide binding protein quorum sensing system (Fontaine et al., 2010; Gardan et al., 2009). These are known to regulate biofilm formation and virulence (Hammer & Bassler, 2003; Kong et al., 2006; Preda & Săndulescu, 2019). The strong signal in carbon metabolism and oxidative phosphorylation were likely reflective of the variation in anaerobic bacteria, which accumulate during gingivitis (Chen et al., 2019; Diaz et al., 2002; Loesche, 1969). An interesting link between salivary metabolites and microbiome functional potential may have been observed in the nucleoside/purine/pyrimidine biosynthesis pathways: while urea was one of the best modelled salivary metabolites and is a potential biomarker for oral health (D’souza et al., 2023; Gaál Kovalčíková et al., 2019; Nascimento et al., 2009; Osmani, 2023), a role of salivary nucleosides in plaque formation or gingivitis is as yet unknown. While many of these functions can be identified as housekeeping activities, we protected ourselves from wrongfully identifying such generic activities as enriched by (1) using SetRank to correct for multiple pathway membership and (2) creating a custom database of pathway elements and removing very large pathways that often connect to such activities. Further research is needed to investigate the relevance of these enriched pathways in the context of dental plaque formation and gingivitis.

In conclusion, we show that PARAFAC modelling of longitudinal oral microbiome and salivary metabolomics data can be used to identify clinically relevant variation and similarly responding microbial subcommunities related to gingivitis. By performing high level integration of predicted microbiome functions and the salivary metabolome, we highlight biochemical pathways of oral biofilm formation and maturation with a likely role in gingivitis.

## Methods

### Data description

The gingivitis challenge study was carried out at the Academic Centre for Dentistry Amsterdam (ACTA) with 15 systemically healthy males and 26 systemically healthy females between the ages of 18 and 55. Recruitment details and exclusion criteria were previously described (Prodan et al., 2016). Reversible gingivitis was induced by omission of oral hygiene during the two-week gingivitis challenge (day 0 to day 14), which was preceded by a two-week baseline period (day −14 to day 0) and followed by a one-week resolution phase (day 14 to day 21; **Figure 1A**). Assessment of plaque formation, bleeding and the collection of oral samples for microbiome were performed throughout the baseline, challenge and resolution phases. Plaque and bleeding were assessed clinically in a half mouth randomized contralateral model (Bentley & Disney, 1995). Oral samples for microbiome analysis were taken at six sites in the mouth: supragingival plaque at the lower and upper jaw lingual surfaces, supragingival plaque at the lower and upper jaw interproximal surfaces, tongue and saliva. The salivary metabolome was only sampled during the challenge phase.

### Red fluorescence imaging

Acquisition of red fluorescent plaque (RF) photographs has been described previously (Heinrich-Weltzien et al., 2003; van der Veen et al., 2016; Volgenant et al., 2017). Briefly, fluorescence photographs were taken of the vestibular aspect of the anterior teeth (cuspid to cuspid, upper and lower jaw) in end-to-end position at every time point using a QLF-D camera (Inspektor Research Systems BV, Amsterdam, the Netherlands). The photographs were assessed planimetrically for the percentage red fluorescent protein (RFP) coverage using RFP analysis software (QA2 V1.25, Inspektor Research Systems BV, Amsterdam, the Netherlands). Three response groups were determined using the RF% values on day 14: low (0-1.7%), mid (1.8-6.0%) and high (6.0%+) (**Figure 1B**) (van der Veen et al., 2016).

### DNA isolation and 16S rRNA gene amplicon sequencing

DNA was isolated from the samples using a previously described method (Zaura et al., 2017). Samples were mixed with 300 µl lysis buffer (Agowa, Berlin, Germany), 500 µl phenol saturated with tris-HCl (pH 8.0) and 500 µl zirconium beads (0.1 mm; BioSpec Products, Bartlesville, OK, USA), and shaken in a bead beater for 3 min at 2800 oscillations/min. DNA released was purified using magnetic beads (Agowa, Berlin, Germany) and used for amplicon sequencing.

The V4 hypervariable region of the 16S rRNA gene was targeted using primers F515 (5’-GTG CCA GCM GCC GCG GTA A-3’) and R806 (5’-GGA CTA CHV GGG TWT CTA AT-3’). The primers included Illumina adapters and a unique 8-nucleotide sample index sequence key (Kozich et al., 2013). PCR was performed using the Phusion Hot Start II High Fidelity PCR Master Mix (Thermo Scientific, Waltham, MA, USA) with 100 pg template. The following amplification program was used: initial denaturation at 98°C for 30 s; 30 cycles of 98°C for 10 s, 55°C for 30 s and 72°C for 30 s; and final elongation at 72°C for 5 min. The amplicon libraries were pooled in equimolar amounts and purified using the QIAquick Gel Extraction Kit (Qiagen, Valencia, CA, USA). Amplicon quality and size were analysed on a Fragment Analyzer (Advanced Analytical Technologies Inc., Ankeny, IA, USA). Amplicon sequencing was performed on the Illumina MiSeq platform (Illumina Technologies, San Diego, CA, USA) using 2 x 200 cycle paired-end settings.

### 16S rRNA gene amplicon data pre-processing

The 16S rRNA gene sequencing data were pre-processed using DADA2 (version 1.14.0; (Callahan et al., 2016)). All microbiome sampling locations except for saliva were pre-processed with default DADA2 settings (truncLen = c(100, 100), maxN = 0, maxEE = c(2,2), truncQ = 2, rm.phix = TRUE, compress = FALSE, matchIDs = TRUE, multithread = TRUE). The saliva samples were processed with a shorter trimming setting (truncLen = c(200, 215)) due to lower quality reads. Taxonomic classification of ASVs was performed using the SILVA NR99 (v138) and HOMD (v15.22) reference databases.

### Metabolomics sampling and profiling

Saliva sample collection and metabolite profiling were described previously (Prodan et al., 2016). Unstimulated saliva was collected at the intervention time points. Participants were instructed to allow saliva to accumulate on the floor of the mouth and to spit at 30 second intervals into a pre-weighted 30 ml polypropylene tube. The collection period was 5 minutes. Non-targeted metabolite profiling of the saliva samples for the gingivitis challenge time points was performed by Metabolon. Samples were processed as described in Metabolon’s standard method (Evans et al., 2009). Raw data were extracted, peak-identified and quality controlled using Metabolon’s hardware and software. Compounds were identified by comparison with Metabolon’s reference library (Lawton et al., 2008). Metabolites were annotated with KEGG compound IDs (Kanehisa et al., 2021; Kanehisa & Goto, 2000).

### Assessment of clinical parameters of gingivitis

Plaque and bleeding scores were clinically assessed by two experts in a half mouth randomized contralateral model, as described previously (van der Veen et al., 2016). Plaque was quantified using a modified Silness & Loë Plaque Index at six sites of the buccal and lingual aspects of all present teeth (Silness & Löe, 1964). Gingival bleeding was quantified using the bleeding on marginal probing index on six gingival areas on the buccal and lingual sides of all present teeth (Van der Weijden et al., 1993).

### Statistical Analysis - Microbiome data processing

The ASV data and taxonomic information were processed using MATLAB (version 2022a). The data were separated by oral site. To limit the sparsity per dataset, ASVs were kept if the sparsity was <50% in any response group (**Supplementary Figure A6**). All other ASVs were removed from the data. By setting a response group-based sparsity-cutoff per ASV, we avoid removing biologically relevant ASVs that would occur in only one or two response groups. ASVs were also removed if they corresponded to chloroplast or mitochondrial sequences, as these were not relevant for the study. An overview of the number of ASVs before and after filtering per sample type is reported in **Supplementary Table B5**. Subsequently, a centered-log ratio transformation - using a pseudo-count of 1 - was performed to correct for compositionality (Aitchison, 1982; Gloor et al., 2017). The datasets were then converted to three-way arrays (**Figure 1D**). Missing samples were kept in the data cube as a row of missing values (**Supplementary Table B6**). This is possible because PARAFAC interpolates the missing data automatically in its alternating least-squares algorithm and maximises the amount of information for the modelling procedure as the other samples of the subject do not have to be removed entirely. Subsequently, centering across the subject mode and scaling within the ASV mode - ignoring missing values - was performed to make the samples comparable at every timepoint and the ASVs comparable for all time points (Bro & Smilde, 2003).

### Statistical Analysis - Salivary metabolomics data processing

The salivary metabolomics data were processed using MATLAB (version 2022a). To limit the sparsity per dataset, metabolites were excluded if they contained more than 25% values below the detection limit (**Supplementary Figure A7**). Additionally, xenobiotic compounds were manually assessed for occurrence across response groups and selected if they were prevalent in most subjects (**Supplementary Data**). After feature selection, 400 of the 499 metabolites remained (**Supplementary Table B5**). Values below the detection limit were imputed with a random value between 0 and the detection limit per metabolite to preserve their distribution. To correct for the dilution caused by the amount of water in the saliva samples, Probabilistic Quotient Normalization (PQN) was performed using the median value of each metabolite as an artificial reference sample (Dieterle et al., 2006). Next, the dataset was (natural) log transformed to stabilize the variance. The dataset was then converted to a three-way array. Subsequently, centering across the subject mode and scaling within the ASV mode - ignoring missing values - was performed to make the samples comparable at every timepoint and the ASVs comparable for all time points (Bro & Smilde, 2003). All subjects were fully sampled, except for subject 3CN8CB for whom no metabolome data was available (**Supplementary Table B6**).

### Functional profile prediction of microbiome data

A functional profile prediction based on the microbiome ASV count data was performed using the Tax4Fun2 package in R (Tax4Fun2 version 1.1.5, R version 4.0.3) (Aßhauer et al., 2015; Wemheuer et al., 2020). ASVs corresponding to Chloroplast or Mitochondria were removed. Samples were rarefied to 10 000 reads to make them comparable (**Supplementary Figures A8** and **A9**). Samples with fewer total reads were removed. This step removed 53 of 1 692 samples. ASVs without counts after rarefying were removed. All steps together removed 6 903 out of 14 014 ASVs. Tax4Fun2 was run with default settings against the included Ref99NR database (Wemheuer et al., 2020) and a custom database based on the HOMD genomes (v9.14, accessed 2020-11-09) (Chen et al., 2010). Molecular functions of the genomes were assigned using Tax4Fun2’s DIAMOND wrapper (Buchfink et al., 2015). HOMD-annotated 16S rRNA gene sequences were extracted from the genomes, cut to the 515F-806R fragment used in this study using cutadapt v1.18 (Martin, 2011). Exact duplication within genomes were removed, and 16S rRNA gene sequence fragments were combined with the functional assignments using Tax4Fun’s generateUserDataByClustering function. The resulting functional prediction profiles - which contain values between 0 and 1 for each function - were used for subsequent analysis. The fraction of unused taxonomic units and the fraction of unused sequences are reported per sample (**Supplementary Figures A10** and **A11**).

### Statistical Analysis - Processing of functional profile predictions

The functionally predicted microbiome data were processed in MATLAB (version 2022a). The data were separated by sample type. To make the data comparable to the salivary metabolomics data, predicted KEGG orthologous groups (KOs) were removed if they did not belong to pathways that the metabolites mapped to. This was done by mapping the KOs and metabolite compound IDs to pathways using the KEGG API (Kawashima et al., 2003). To limit the sparsity per dataset, KOs were removed if the number of zeroes for all response groups was >50% (**Supplementary Figure A12**). Additionally, KOs were removed if the sum-of-squares was lower than 0.025% of the total sum-of-squares of the dataset (**Supplementary Figure A13**). This latter filter was to ensure that the modelling procedure focused on relevant variation in the data and made the total number of variables comparable to the metabolomics data, while retaining most information. An overview of the number of KOs remaining after feature selection is reported (**Supplementary Table B5**). Subsequently a centered-log ratio transformation of each dataset was performed to correct for compositionality (Aitchison, 1982; Gloor et al., 2017). The dataset was then converted to a three-way array. Missing samples were kept in the data cube as a row of missing values (**Supplementary Table B6**). Subsequently, centering across the subject mode and scaling within the ASV mode - ignoring missing values - was performed to make the samples comparable at every timepoint and the ASVs comparable for all time points (Bro & Smilde, 2003).

### Statistical Analysis - Parallel Factor Analysis (PARAFAC)

Details on the creation of PARAFAC models for microbiome (Martino et al., 2021) and metabolomics data (Bro, 1997) have been described elsewhere. The PARAFAC implementation from the N-way toolbox (version 1.8.0.0) in MATLAB (version 2022a) was used to create PARAFAC models for all datasets (Bro, accessed 26-10-2023). Similar to PCA, the correct number of components of the PARAFAC model needed to be determined to create an optimal model (Bro & Kiers, 2003; van der Ploeg et al., 2024). This was done by inspecting the number of iterations needed to converge (Bro, 1997), the core consistency diagnostic (CORCONDIA) (Bro & Kiers, 2003), the variation explained (Bro, 1997), and the Tucker congruence coefficient per mode (Lorenzo-Seva & ten Berge, 2006; Tucker, 1951). Additionally, a jack-knife approach was used to determine the stability of the PARAFAC modelling procedure. These metrics were inspected per component for every generated model (**Supplementary Data**). All generated models are reported in **Supplementary Figures A14-A26**. An overview of the model statistics also reported (**Supplementary Table B7**). Congruence loadings, which describe the relationships between the original variables of the dataset and the latent variables of the corresponding model, were calculated for every component in every sample type (Lorho et al., 2006). While the metrics above suggested a one-component PARAFAC model corresponding to the microbiome data at the lower jaw interproximal niche, the model described less than 10% of the variation in the data. Furthermore, manual inspection revealed that the model did not describe variation that we could explain biologically. Hence, we did not analyse this model further in subsequent steps.

A transformation of the subject loadings was required for comparison with the longitudinally measured clinical parameters of gingivitis. The ASV loading vectors were orthonormalized using the Gram-Schmidt orthonormalization procedure in the pracma R package (version 2.4.2) (Borchers & Borchers, 2022; Schmidt, 1989). The (same) transformation matrix is then applied to the Kronecker product of the subject and the time loadings to obtain interpretable loadings of every subject-time combination (Kiers, 2000). The correlation of the time-resolved subject loadings with the clinical parameters of gingivitis was then tested (**Table 1** and **Supplementary Table B1**).

### Statistical Analysis - Microbiome ASV cluster analysis

Loading plots of the ASV mode and the subject mode for each of the microbiome PARAFAC models were created (**Figure 2**). ASVs were filtered out if they had a variation explained lower than the average of the model or a congruence loading lower than 0.4 (Lorho et al., 2006). ASVs were clustered based on their fitted abundances according to the PARAFAC model using the K-medoids algorithm from the cluster R package (version 2.1.4) with 50 random starts to be robust against outliers (Maechler, 2019). The number of clusters was determined using the within-cluster sum of squares, silhouette width and gap statistic metrics as reported by the factoextra R package (version 1.0.7, **Supplementary Figure A27**) (Kassambara & Mundt, 2017). An overview of the taxonomic information per cluster can be found in **Table 2**. Bacteria in **Table 2** were only reported if two criteria were met: (1) the taxonomic classification was resolved to species level by DADA2 using the SILVA (v138) and HOMD (v15.22) databases and (2) the annotation agreed with a separate best hit annotation run using the HOMD (v15.22) database. Cases where the classification was resolved to species level by the DADA2 run but only to the genus level by the separate HOMD run are also reported.

The response to the gingivitis intervention of the ASV clusters was shown using the relative abundance sum per ASV cluster derived from the original count data (**Figure 3** and **Supplementary Figure A3**). The mean of the summed relative abundances and standard error of the mean were calculated per response group for a given sample type and time point. The mean difference between the high and low responders was tested using a permutation analysis where the response group membership of the subjects was permuted (**Supplementary Table B2**).

### Statistical Analysis - Functional enrichment analysis

The PARAFAC models of the functionally profiled microbiome and salivary metabolomics data were combined to perform a pathway enrichment test per sample type. This was done by calculating the variance explained and congruence loadings per KO or metabolite in each model (Lorho et al., 2006). If the PARAFAC model contained two components, the maximum of the two congruence loadings per feature was used. The variance explained and congruences were normalized by the maximum value per model prior to integration. The normalized variance explained and normalized congruence per feature were averaged to obtain a ranking of pathway elements (KOs and metabolites). This causes the best modelled KO and compound ID to be at the top of this list. The ranked pathway element list was cut off at 75% of its length to ensure a good separation between well-modelled and averagely-modelled pathway elements (**Supplementary Figure A28**).

The SetRank R package (version 1.1.0) was used to obtain pathways of interest (Simillion et al., 2017). SetRank is a gene set enrichment algorithm that obtains a high sensitivity by correcting for multiple pathway membership. A custom database of pathway elements was created by combining the KO and compound ID to pathway mappings obtained through the KEGG API (**Supplementary Data**). Pathways were removed from our database if they could not be performed by prokaryotes. Generic, large pathways with more than 750 elements were removed to ensure that the enrichment results were specific enough for interpretation (**Supplementary Figure A29**). Subsequently the SetRank analysis was performed with standard settings per sample type. The multiple-pathway corrected p-values are reported (p≤0.05, **Figure 4** and **Supplementary Table B4**)

## Data Availability

The ASV abundance data is available at https://github.com/GRvanderPloeg/TIFN-multiway. Raw sequencing data is available by request from the authors, in accordance with the informed consent signed by the study participants.

## Code Availability

The underlying code for this study is available on GitHub and can be accessed via https://github.com/GRvanderPloeg/TIFN-multiway/releases/tag/v1.2.

## Ethics statement

The study involving human participants was conducted in accordance with the ethical principles of the 64th WMA Declaration of Helsinki (October 2013, Brazil) and the Medical Research Involving Human Subjects Act (WMO), approximating Good clinical Practice (CPMP/ICH/135/95) guidelines. The clinical trial was approved by the Medical Ethical Committee of the VU Medical Center (2014.505) and registered at the public trial register of the Central Committee on Research Involving Human Subjects (CCMO) under number NL51111.029.14.

## Author Contributions

G.R. van der Ploeg: formal analysis, methodology, writing - original draft; B.W. Brandt: investigation, writing - review & editing; B.J.F. Keijser: investigation, project administration, writing - review & editing; M.H. van der Veen: investigation, project administration, writing - review & editing; C.M.C. Volgenant: investigation, writing - review & editing; E. Zaura: conceptualization, investigation, project administration, funding acquisition, writing - review & editing; A.K. Smilde: conceptualization, methodology, supervision, funding acquisition, writing - review & editing; J.A. Westerhuis: conceptualization, methodology, supervision, formal analysis, writing - original draft; A. Heintz-Buschart: conceptualization, formal analysis, supervision, project administration, writing - original draft. All authors reviewed and approved the manuscript.

## Competing interests

The authors declare that the research was conducted in the absence of any commercial or financial relationships that could be construed as a potential conflict of interest.

## Acknowledgements

We want to thank Michelle van der Wurff (TNO) and Tim van den Broek (TNO) for their help in data acquisition, processing and storing and Jesse Alderliesten (UU/UvA), Fred White (UvA), Cynthia Albracht (UvA), Juan Pablo Bascur (CWTS) and Rianne Warmerdam (Aiden) for their useful discussions and suggestions. The authors thank N.A.M. Rosema for coordinating the clinical study. For assistance in capturing fluorescence photographs, the authors thank Y. Altindağ. For the clinical examinations the authors thank J.M. Voll (clinical plaque assessment with Silness & Löe), and S. Bizzarro (assessment of the bleeding on marginal probing).

This research is supported by the Dutch Technology Foundation STW (project number 10948) and the Top Institute Food and Nutrition (TIFN), a public-private partnership on precompetitive research in food and nutrition. Organizations supporting this project had no role in study design, data collection and analysis, decision to publish, or preparation of the manuscript. QLF data was obtained with funding from NWO (ZonMw-STW-NIG-program): “Project 10948: Seeing is believing? A novel tool for the visualization of oral disease manifestations.” GRvdP was funded by a grant from the University of Amsterdam, Research Priority Area on Personal Microbiome Health.

## Notes

https://zenodo.org/doi/10.5281/zenodo.10450355

### Competing Interest Statement

The authors have declared no competing interest.

### Summary of Updates

Figure 2 caption revised; Results section on ASV clusters updated to give general description; Methods and Discussion section updated to clarify that only the bacterial microbiome is studied; Discussion and Results sections on SetRank pathway enrichment analysis updated to clarify how we protected ourselves from finding general housekeeping activities as enriched; Table 2 caption and Methods section on Table 2 updated to clarify when clustered species are reported.

