## Supplementary Information for "Multi-way modelling of oral microbial dynamics and host-microbiome interactions during induced gingivitis"

### Appendix A Supplementary Figures

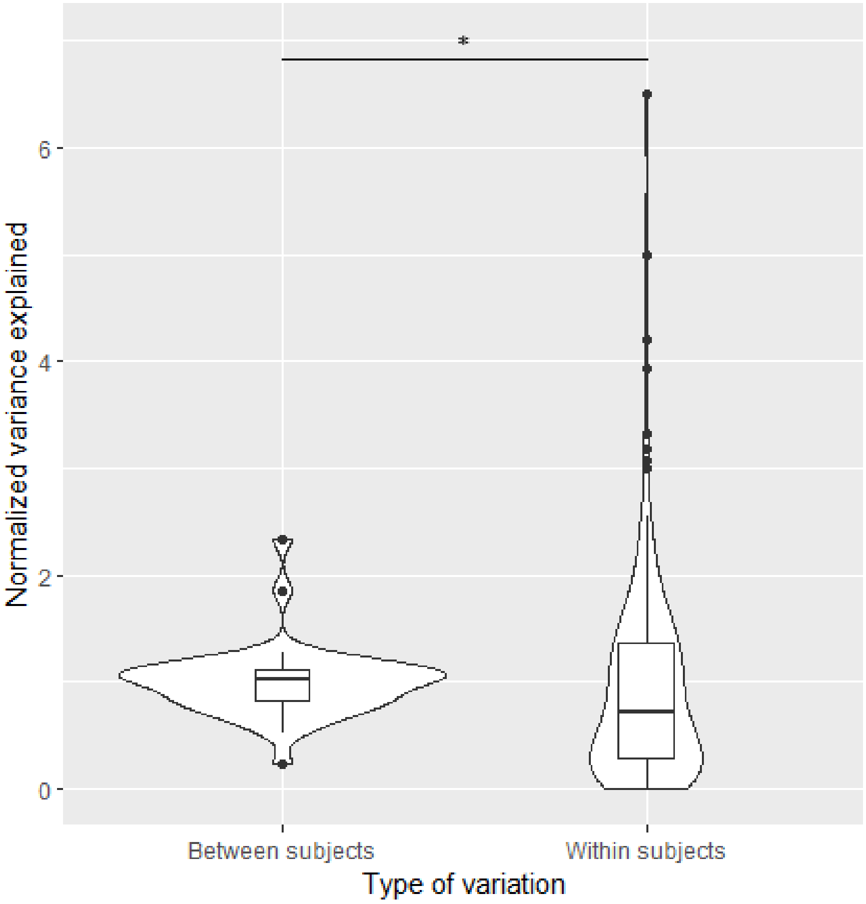


**Fig. A1** Comparison of within (n=246) and between (n=42) subject variation for all microbiome PARAFAC models combined. Modelled variation between subjects was calculated as the variation explained per timepoint in each sampling location. Modelled variation within subjects was calculated as the variation explained per subject over all timepoints in each sampling location. All variation explained values were normalized by the variation of the entire model per sample type to obtain comparable values. (p=0.02; One-sided Wilcoxon rank-sum test)

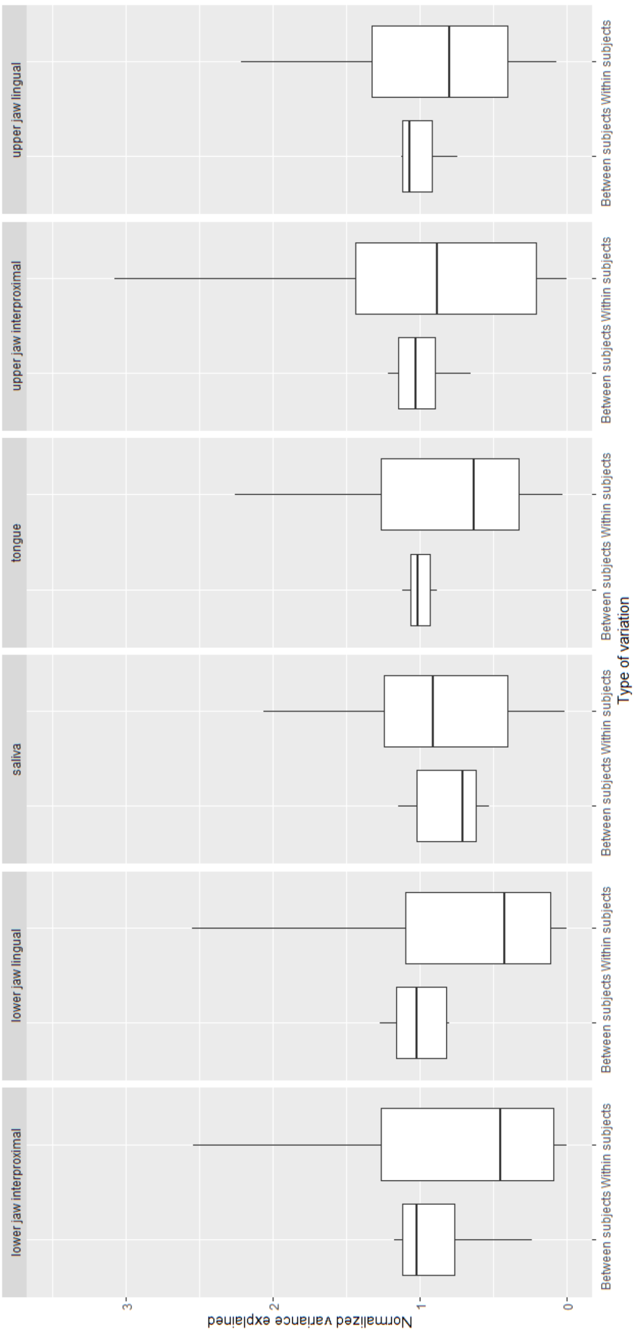


**Fig. A2** Comparison of within (n=41) and between (n=7) subject variation of the PARAFAC models per microbiome sample type. Modelled variation between subjects was calculated as the variation explained per timepoint in each sampling location. Modelled variation within subjects was calculated as the variation explained per subject over all timepoints in each sampling location. Variation explained was normalized by the variation of the entire model per sample type to obtain comparable values. The differences in within and between subject variation per sampling location were not significant (p>0.05; One-sided Wilcoxon rank-sum test).

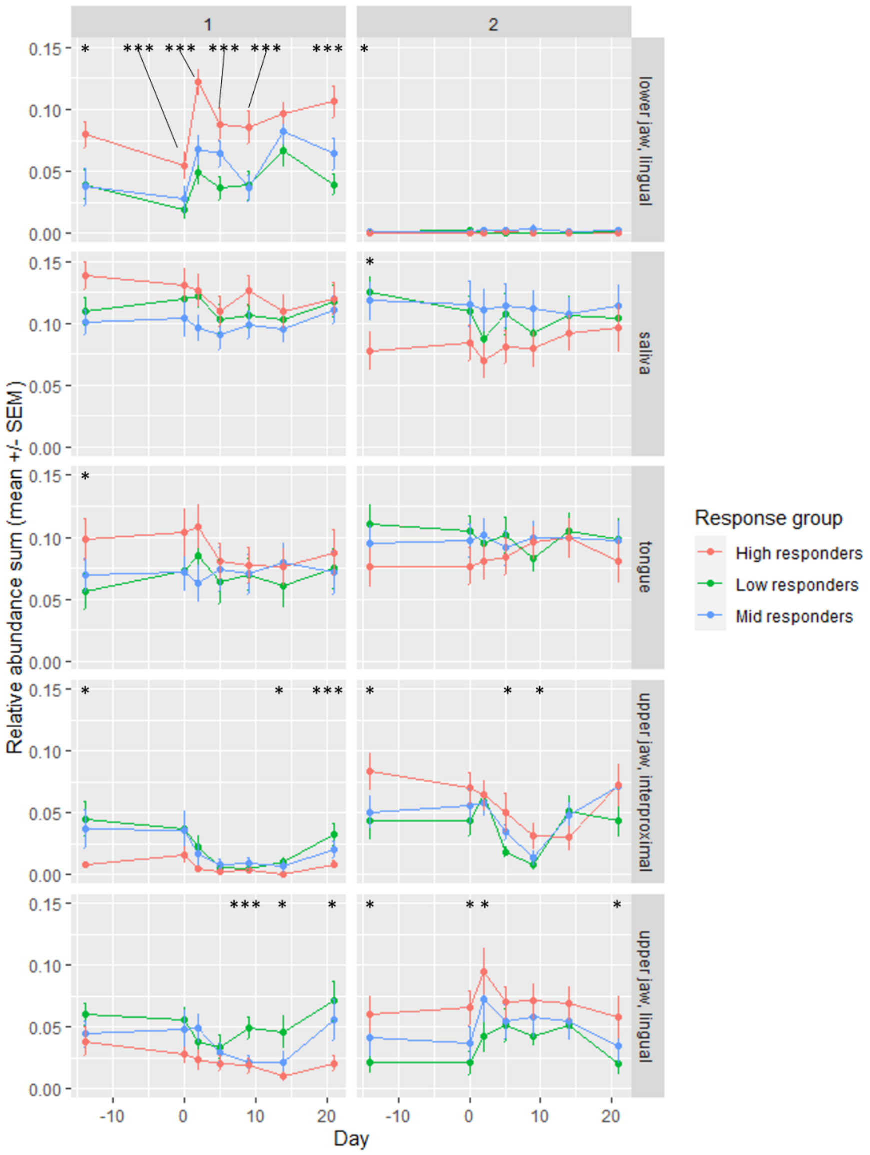


**Fig. A3** Overview of the relative abundance sum per ASV cluster in all sample types, separated by response group. The mean difference between the high and low response group was tested using a permutation test of 999 iterations. P-values were Benjamini-Hochberg corrected. ASV clusters 3 and 4 of the model corresponding to the upper jaw interproximal plaque are not shown for visual clarity. ASV cluster 2 of the model corresponding to the lower jaw lingual samples is not shown here due to very low abundance values (see Figure A4). ASV cluster 1 in the lower jaw lingual samples (containing only *S. salivarius*) has a significantly higher mean relative abundance sum in the high responders compared to the low responders at nearly every time point. Please refer to Table 2 for the list of identified species per ASV cluster. (*: $p \leq 0.05$; **: $p \leq 0.01$; ***: $p \leq 0.001$)

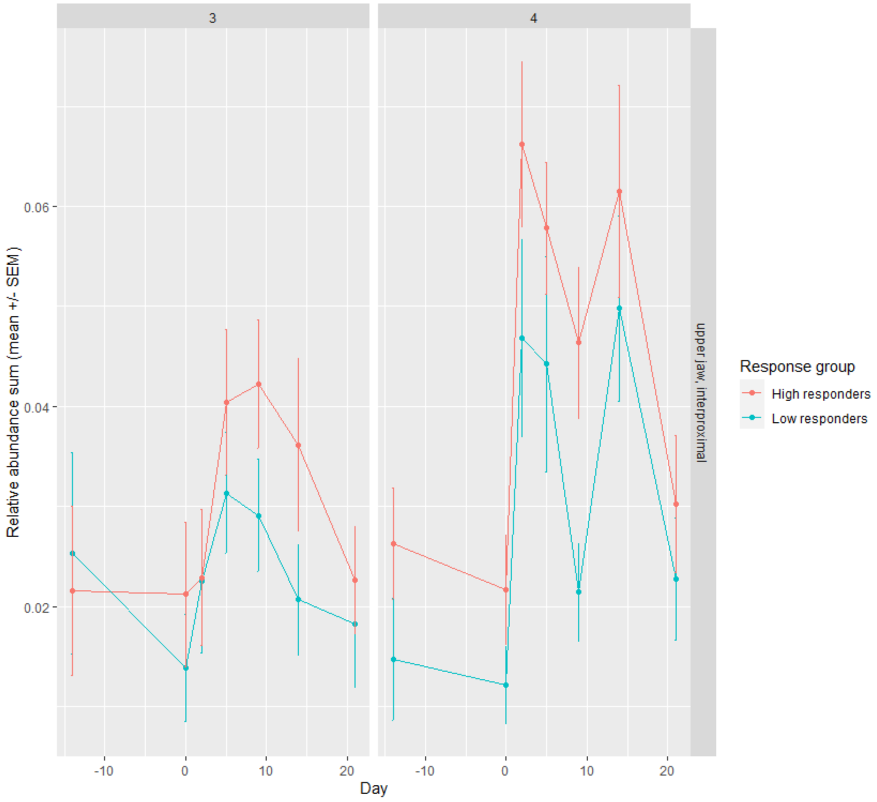


**Fig. A4** Overview of the relative abundance sum for upper jaw interproximal ASV clusters 3 and 4, separated by response group.

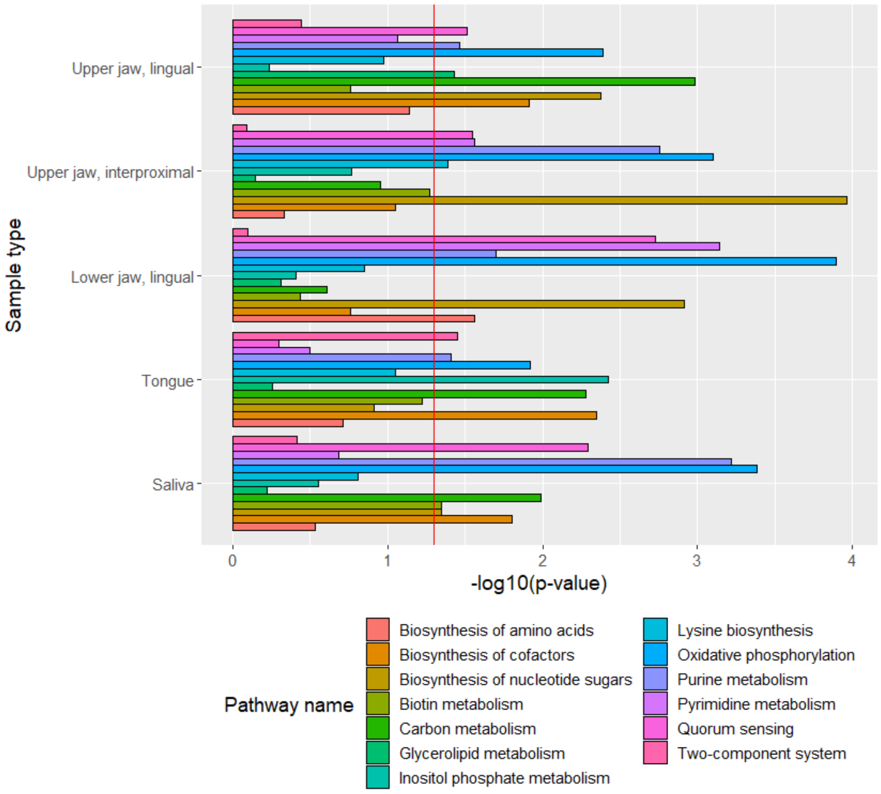


**Fig. A5** Overview of the pathway enrichment results per combination of oral site microbiome and the salivary metabolome. Tax4fun2 was used to create functional predictions from ASV data. Well-modelled metabolites and microbiome molecular functions were integrated by mapping them to KEGG pathway level. SetRank was then used to test for pathway enrichment and corrected for multiple-pathway membership. Pathways here were filtered to be significant ($p \leq 0.05$) in at least one sample type and to contain at least two well-modelled molecular functions and two well-modelled compounds.

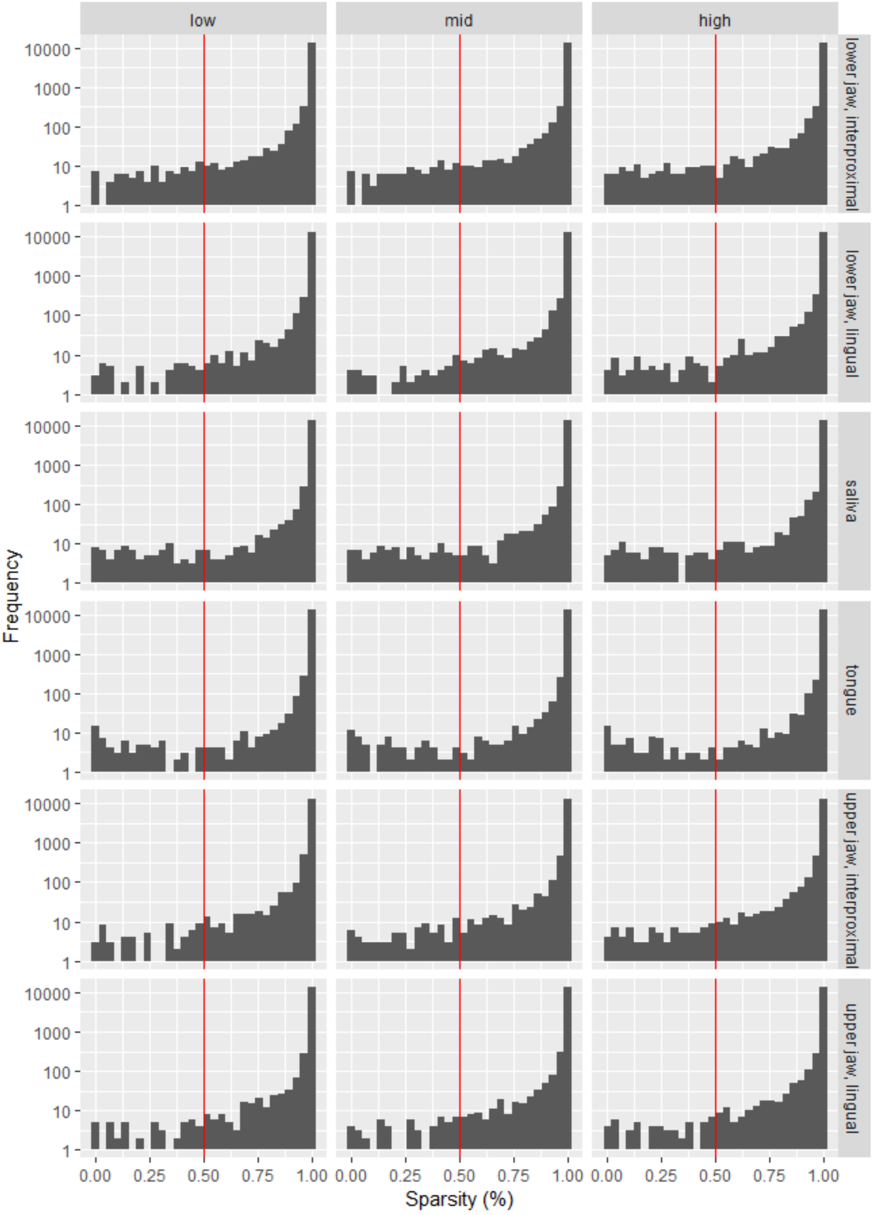


**Fig. A6** Histograms of the sparsity of ASVs per sample type and response group in the microbiome data. Sparsity is here defined as the percentage of measurements of an ASV that are zero. ASVs were removed from the microbiome data if they had a sparsity higher than 0.50 in all response groups, indicated by the red line.

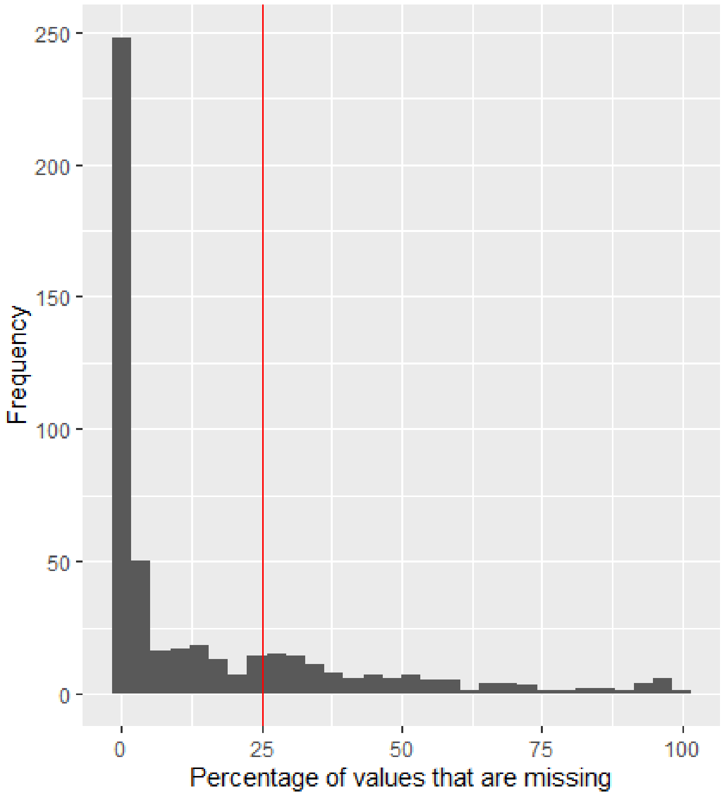


**Fig. A7** Histogram of the percentage missing values per metabolite in the salivary metabolomics data. Values are noted as missing if the amount of the metabolite is below the detection limit. Metabolites were removed from the data if the percentage of missing values was higher than 25%, indicated by the red line.

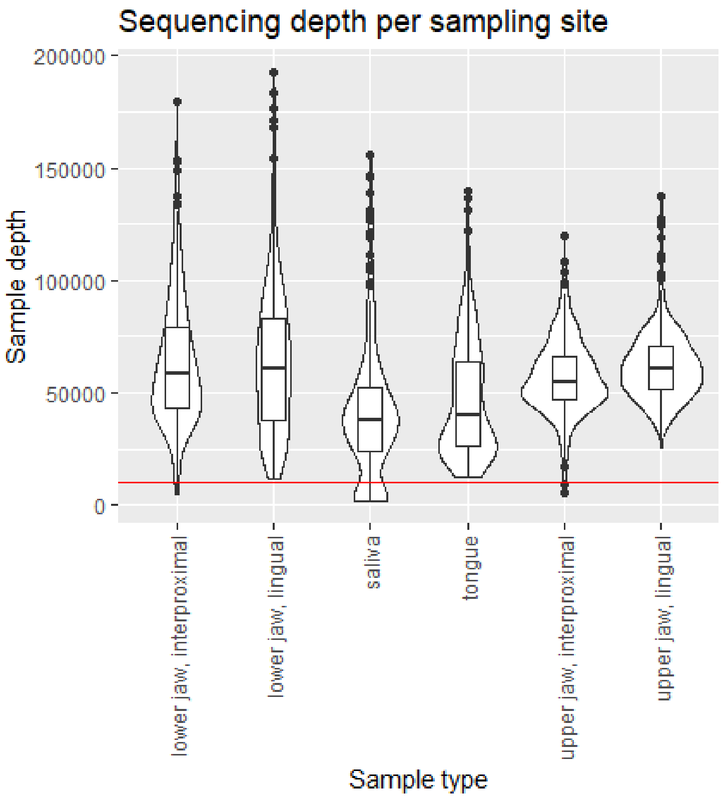


**Fig. A8** Violin plot and boxplot showing the sampling depth per sample type. Whiskers are defined as 1.5 times the interquartile range. The red line at 10.000 reads indicates the rarefaction level chosen for the functional predictions using Tax4Fun2.

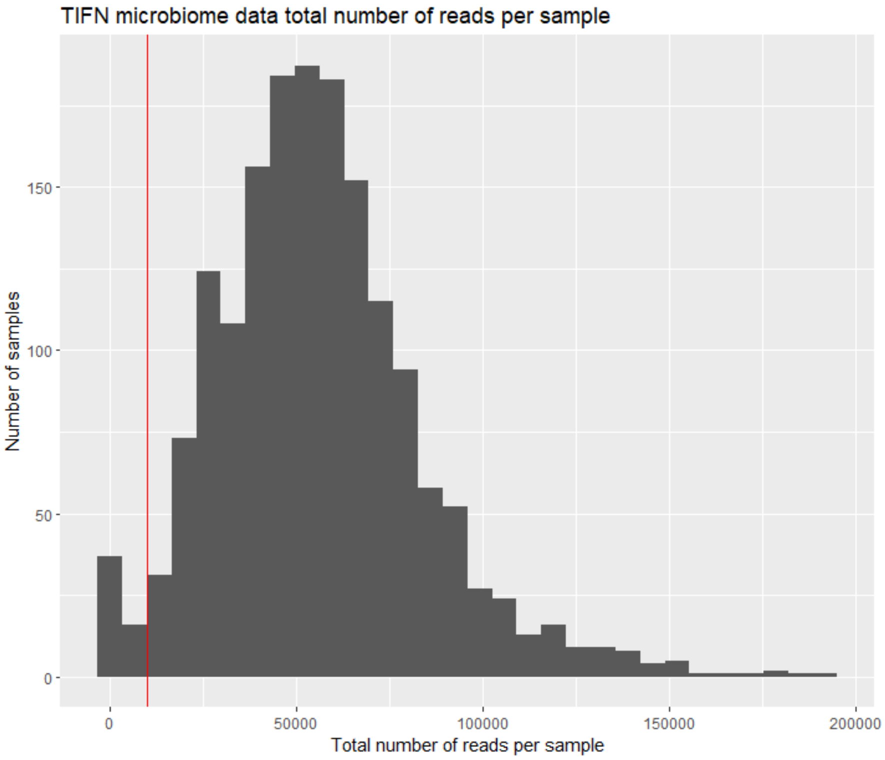


**Fig. A9** Sample depth histogram for all samples in the microbiome data. The red line at 10.000 reads indicated the rarefaction level chosen for the functional predictions using Tax4Fun2.

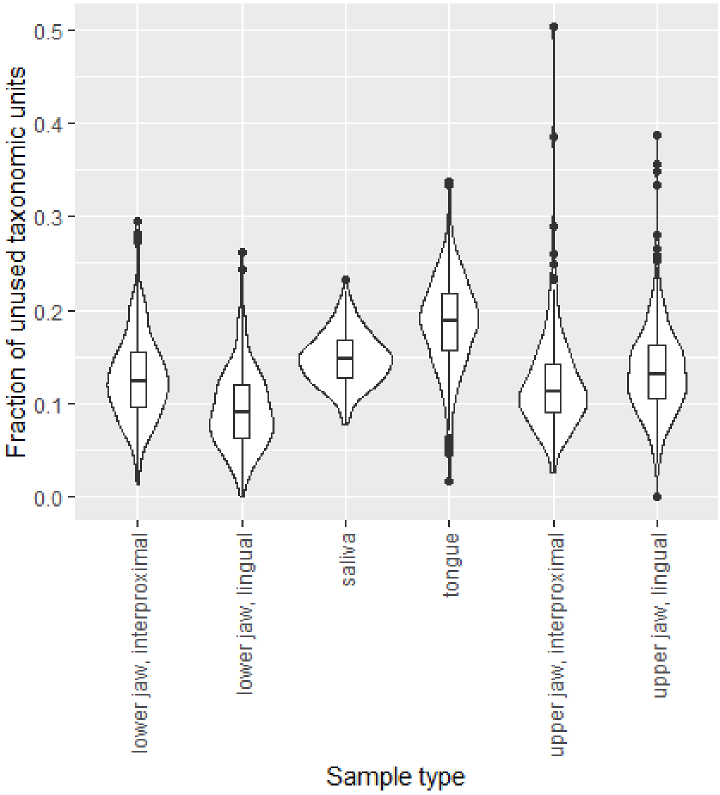


**Fig. A10** Fraction of unused ASVs per sample type as reported in the Tax4Fun2 output. These ASVs have no close hits in the reference data and hence cannot be used for the functional predictions.

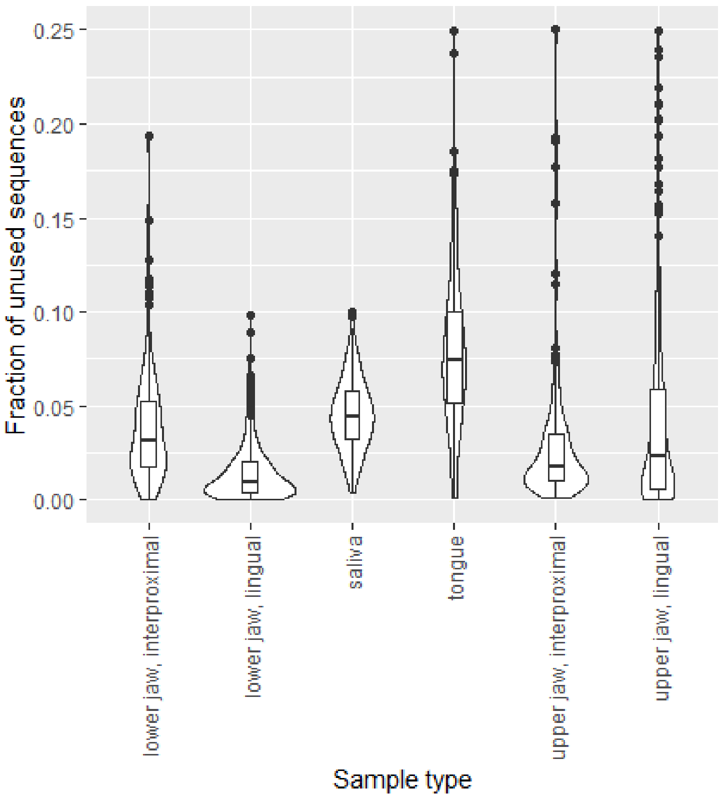


**Fig. A11** Fraction of unused sequences per sample type as reported in the Tax4Fun2 output. These sequences correspond to ASVs that have no close hits in the reference data and hence cannot be used for the functional predictions.

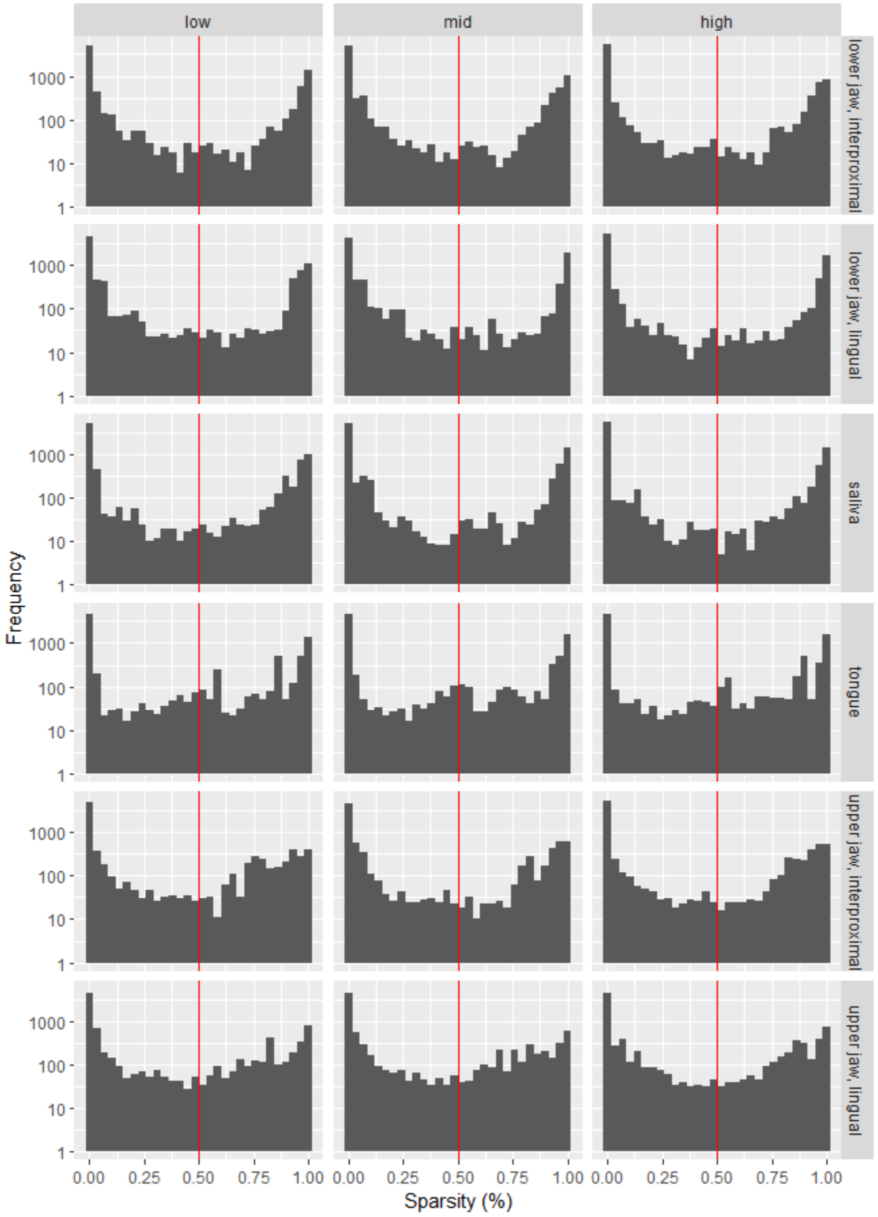


**Fig. A12** Sparsity of functionally predicted KOs per sample type and response group. Sparsity is here defined as the percentage of values per KO that are zero. KOs were removed if they had a sparsity higher than 0.50 in all response groups, indicated by the red line.

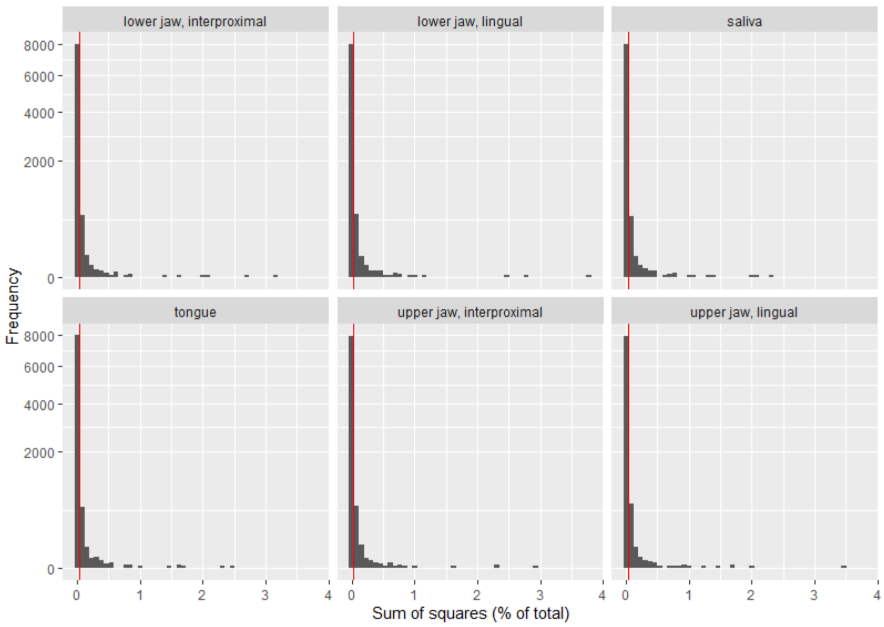


**Fig. A13** Percentage sum-of-squares of functionally predicted KOs per sample type. KOs were removed if they had a percentage sum of squares lower than 0.0025, indicated by the red line.

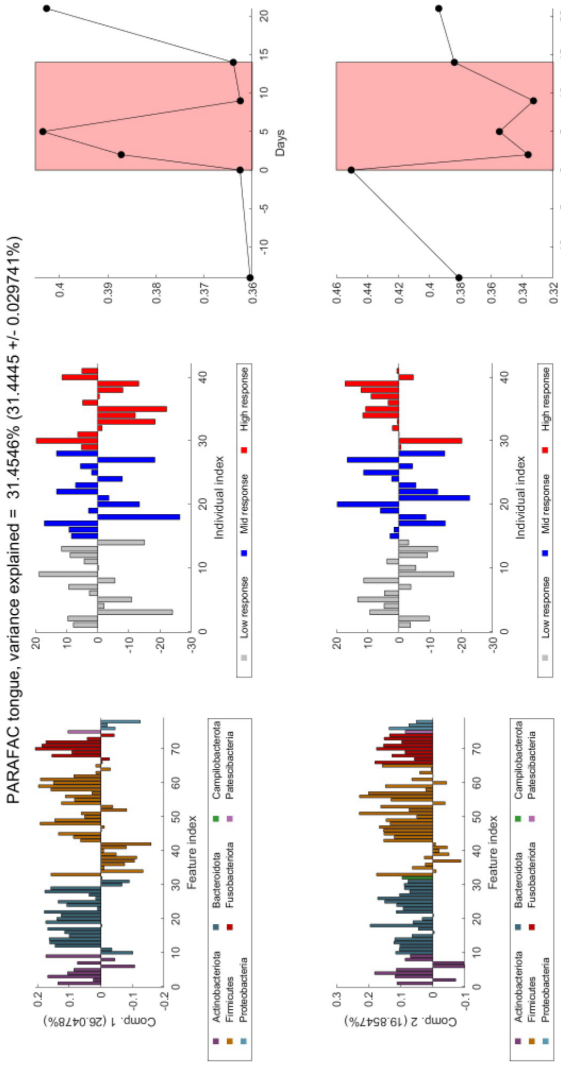


**Fig. A14** Overview of the PARAFAC model corresponding to the samples obtained from the tongue microbiome. This model describes 31.5% of the variation in the data. The first component is shown in the top row and the second component is shown in the bottom row. In the left column, the ASV loadings are shown. In the middle column, the subject loadings are shown. In the right column the time loadings are shown. The ASV loadings are coloured by their phylum level taxonomic classification. The subject loadings are coloured by their response group. In the time loading plot, the gingivitis intervention time points are shown with a red background.

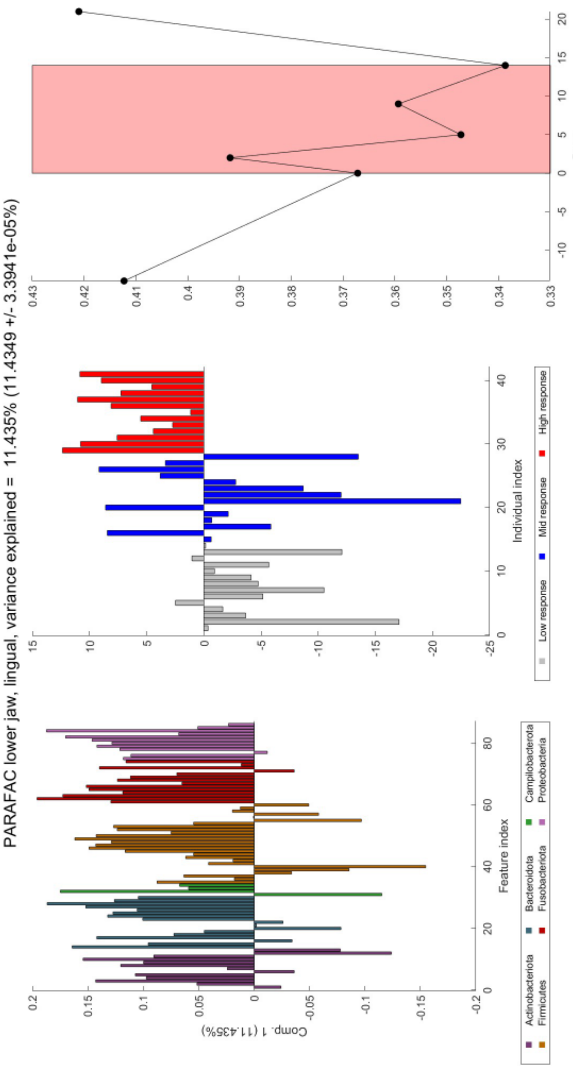


**Fig. A15** Overview of the PARAFAC model corresponding to the samples obtained from the lower jaw lingual microbiome. This model describes 11.4% of the variation in the data. In the left column, the ASV loadings are shown. In the middle column, the subject loadings are shown. In the right column the time loadings are shown. The ASV loadings are coloured by their phylum level taxonomic classification. The subject loadings are coloured by their response group. In the time loading plot, the gingivitis intervention time points are shown with a red background.

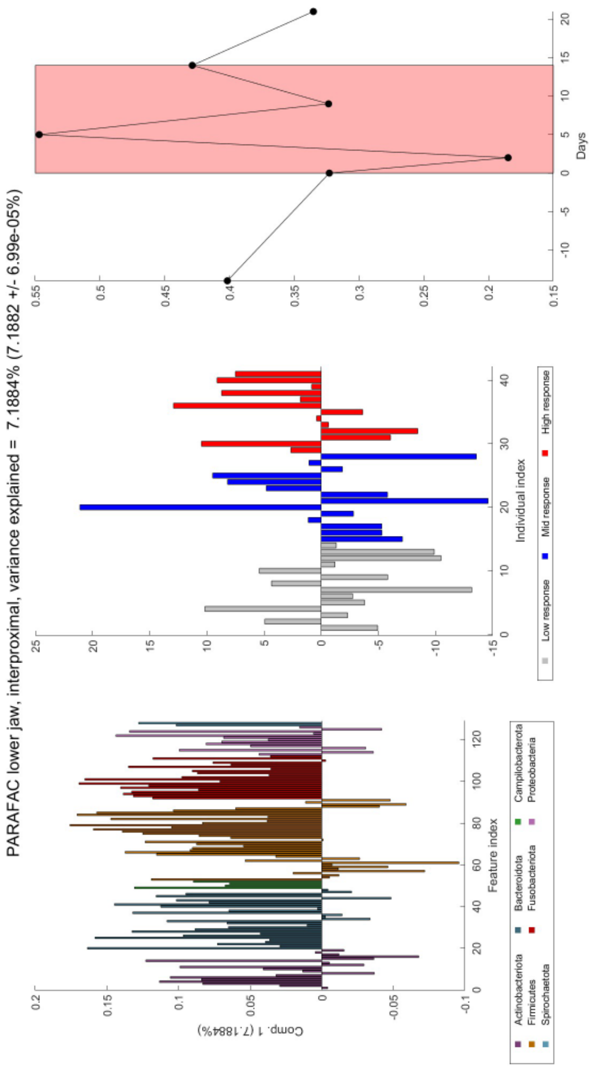


**Fig. A16** Overview of the PARAFAC model corresponding to the samples obtained from the lower jaw inteproximal microbiome. This model describes 7.2% of the variation in the data. In the left column, the ASV loadings are shown. In the middle column, the subject loadings are shown. In the right column the time loadings are shown. The ASV loadings are coloured by their phylum level taxonomic classification. The subject loadings are coloured by their response group. In the time loading plot, the gingivitis intervention time points are shown with a red background.

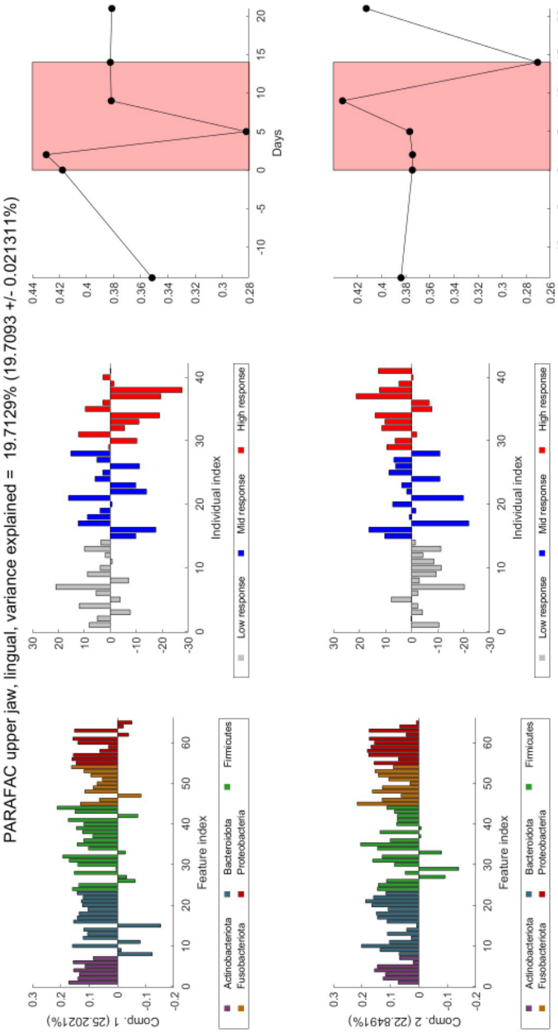


**Fig. A17** Overview of the PARAFAC model corresponding to the samples obtained from the upper jaw lingual microbiome. This model describes 19.7% of the variation in the data. The first component is shown in the top row and the second component is shown in the bottom row. In the left column, the ASV loadings are shown. In the middle column, the subject loadings are shown. In the right column the time loadings are shown. The ASV loadings are coloured by their phylum level taxonomic classification. The subject loadings are coloured by their response group. In the time loading plot, the gingivitis intervention time points are shown with a red background.

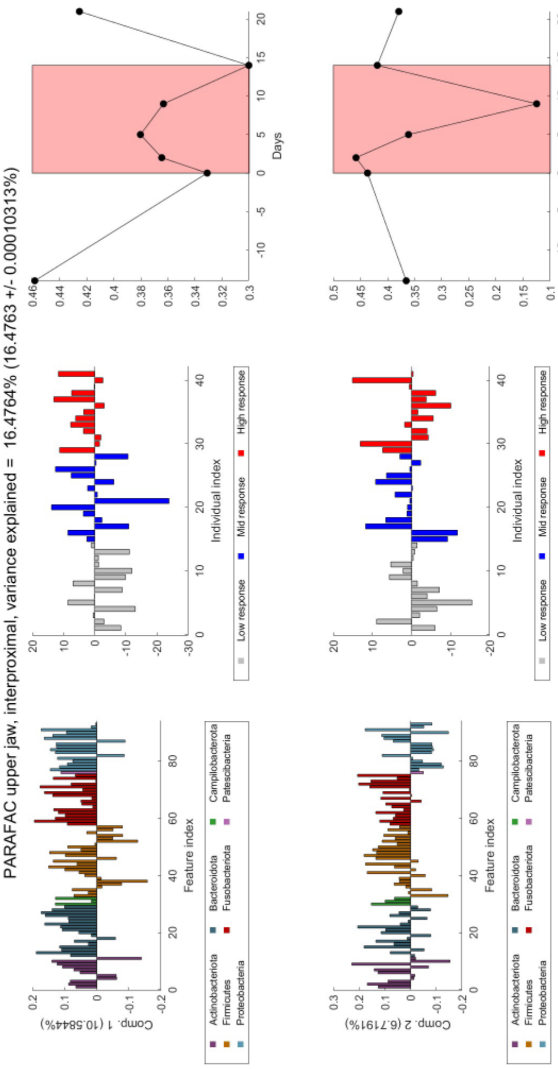


**Fig. A18** Overview of the PARAFAC model corresponding to the samples obtained from the upper jaw interproximal microbiome. This model describes 16.5% of the variation in the data. The first component is shown in the top row and the second component is shown in the bottom row. In the left column, the ASV loadings are shown. In the middle column, the subject loadings are shown. In the right column the time loadings are shown. The ASV loadings are coloured by their phylum level taxonomic classification. The subject loadings are coloured by their response group. In the time loading plot, the gingivitis intervention time points are shown with a red background.

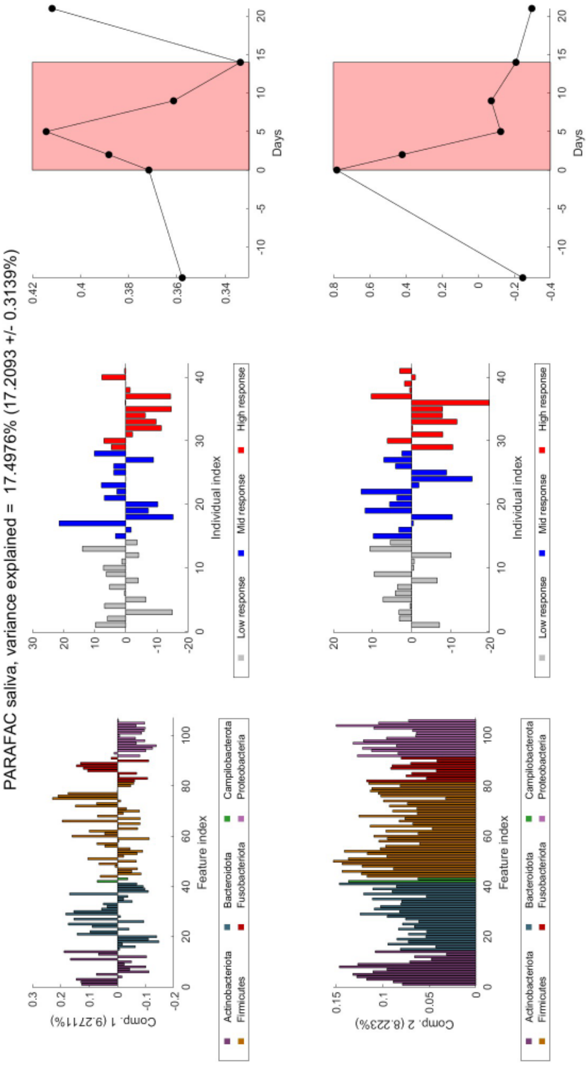


**Fig. A19** Overview of the PARAFAC model corresponding to the samples obtained from the saliva microbiome. This model describes 17.5% of the variation in the data. The first component is shown in the top row and the second component is shown in the bottom row. In the left column, the ASV loadings are shown. In the middle column, the subject loadings are shown. In the right column the time loadings are shown. The ASV loadings are coloured by their phylum level taxonomic classification. The subject loadings are coloured by their response group. In the time loading plot, the gingivitis intervention time points are shown with a red background.

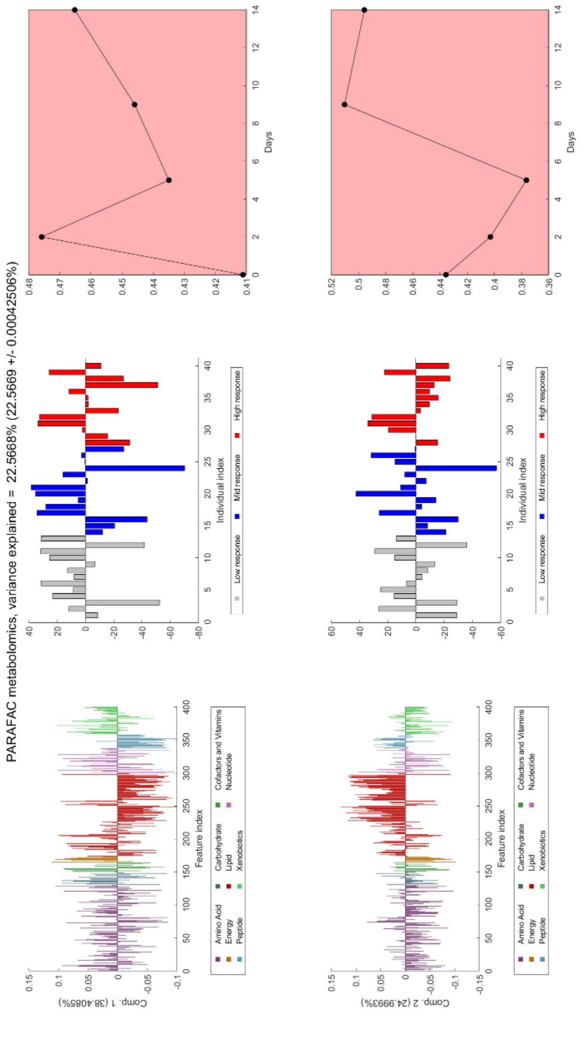


**Fig. A20** Overview of the PARAFAC model corresponding to the samples obtained from the salivary metabolomics. This model describes 22.6% of the variation in the data. The first component is shown in the top row and the second component is shown in the bottom row. In the left column, the metabolite loadings are shown. In the middle column, the subject loadings are shown. In the right column the time loadings are shown. The metabolite loadings are coloured by their metabolite class (see legend). The subject loadings are coloured by their response group. In the time loading plot, the gingivitis intervention time points are shown with a red background. For the salivary metabolomics, only samples taken during the intervention were available.

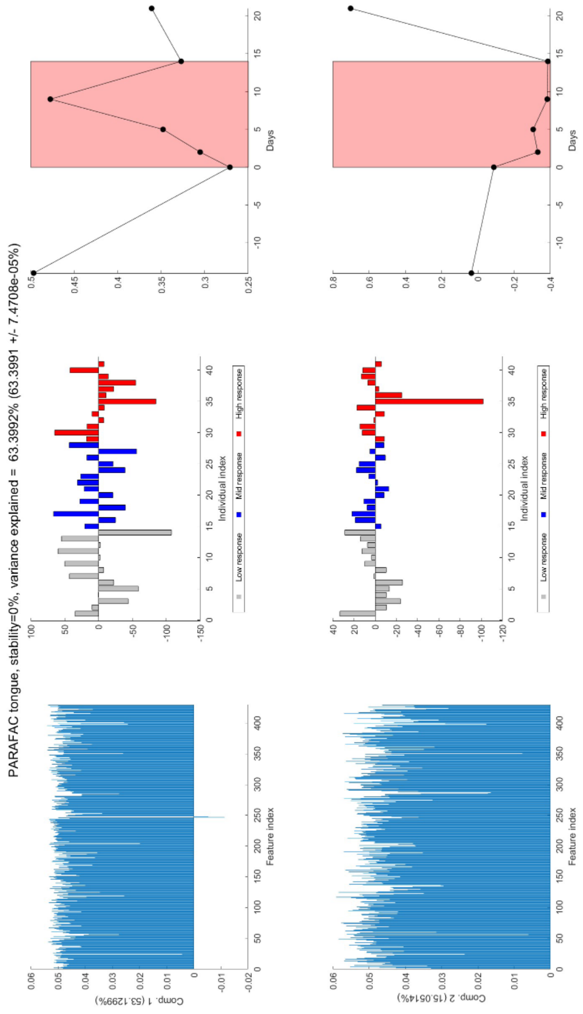


**Fig. A21** Overview of the PARAFAC model corresponding to the functional predictions of the samples obtained from the tongue microbiome data. This model describes 63.4% of the variation in the data. The first component is shown in the top row and the second component is shown in the bottom row. In the left column, the ASV loadings are shown. In the middle column, the subject loadings are shown. In the right column the time loadings are shown. The subject loadings are coloured by their response group. In the time loading plot, the gingivitis intervention time points are shown with a red background.

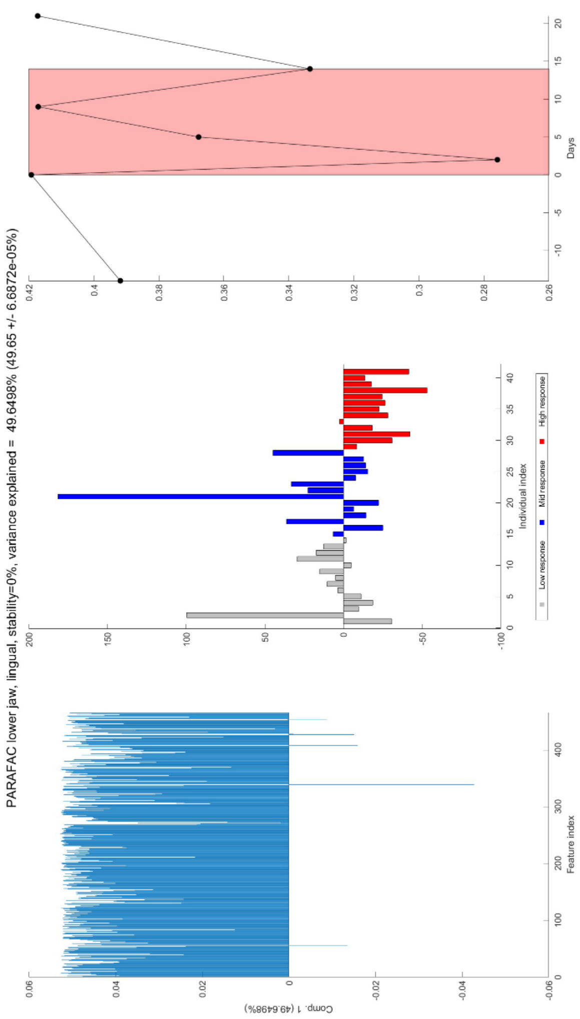


**Fig. A22** Overview of the PARAFAC model corresponding to the functional predictions of the samples obtained from the lower jaw lingual microbiome. This model describes 49.6% of the variation in the data. In the left column, the ASV loadings are shown. In the middle column, the subject loadings are shown. In the right column the time loadings are shown. The subject loadings are coloured by their response group. In the time loading plot, the gingivitis intervention time points are shown with a red background.

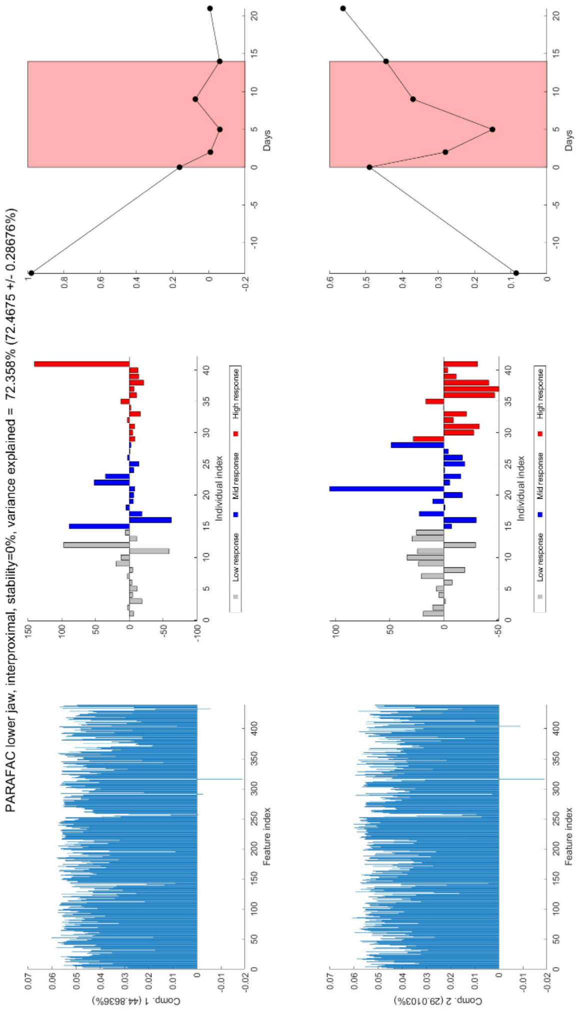


**Fig. A23** Overview of the PARAFAC model corresponding to the functional predictions of the samples obtained from the lower jaw interproximal microbiome. This model describes 72.5% of the variation in the data. The first component is shown in the top row and the second component is shown in the bottom row. In the left column, the ASV loadings are shown. In the middle column, the subject loadings are shown. In the right column the time loadings are shown. The subject loadings are coloured by their response group. In the time loading plot, the gingivitis intervention time points are shown with a red background.

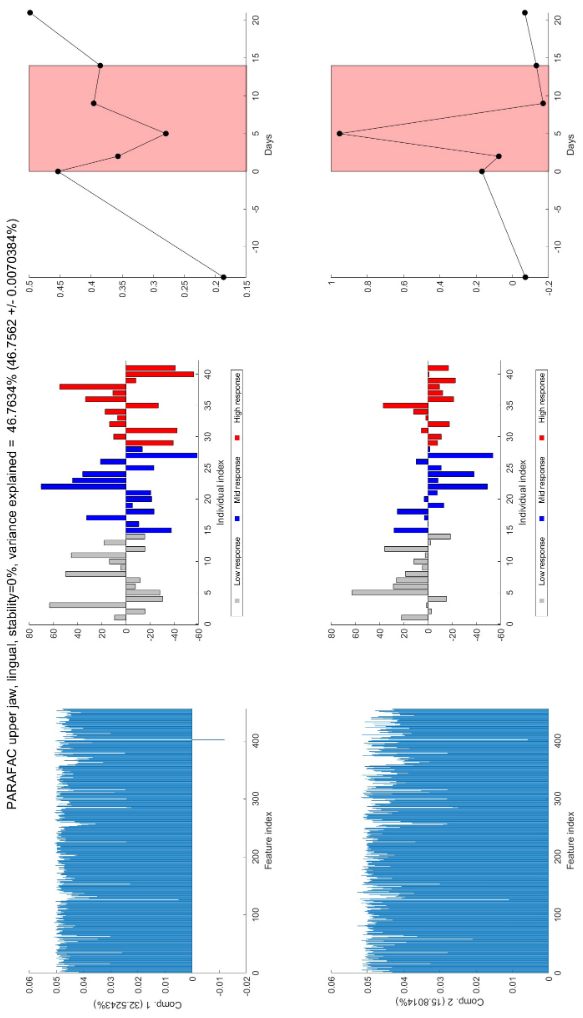


**Fig. A24** Overview of the PARAFAC model corresponding to the functional predictions of the samples obtained from the upper jaw lingual microbiome. This model describes 46.8% of the variation in the data. The first component is shown in the top row and the second component is shown in the bottom row. In the left column, the ASV loadings are shown. In the middle column, the subject loadings are shown. In the right column the time loadings are shown. The subject loadings are coloured by their response group. In the time loading plot, the gingivitis intervention time points are shown with a red background.

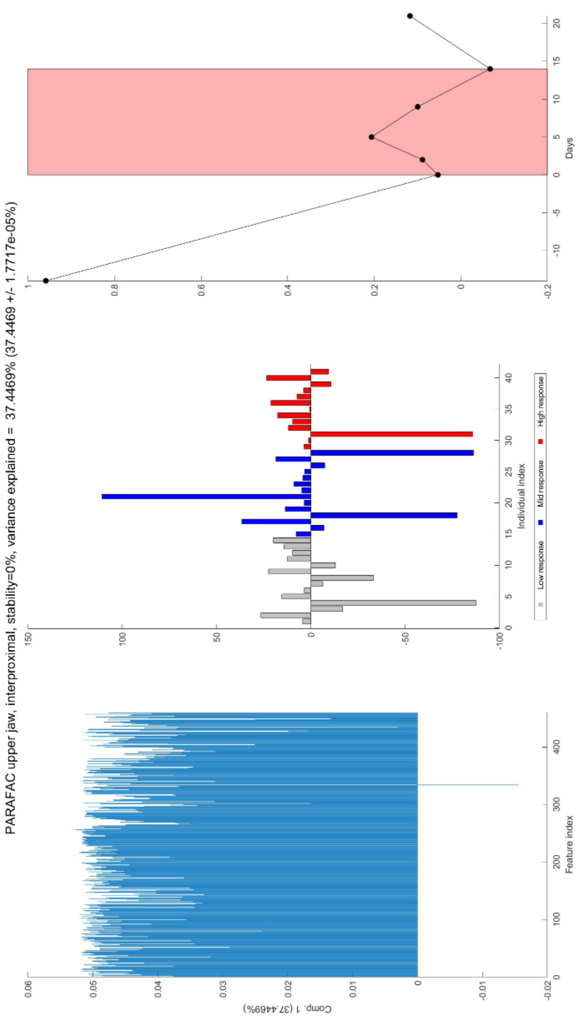


**Fig. A25** Overview of the PARAFAC model corresponding to the functional predictions of the samples obtained from the upper jaw interproximal microbiome. This model describes 37.4% of the variation in the data. In the left column, the ASV loadings are shown. In the middle column, the subject loadings are shown. In the right column the time loadings are shown. The subject loadings are coloured by their response group. In the time loading plot, the gingivitis intervention time points are shown with a red background.

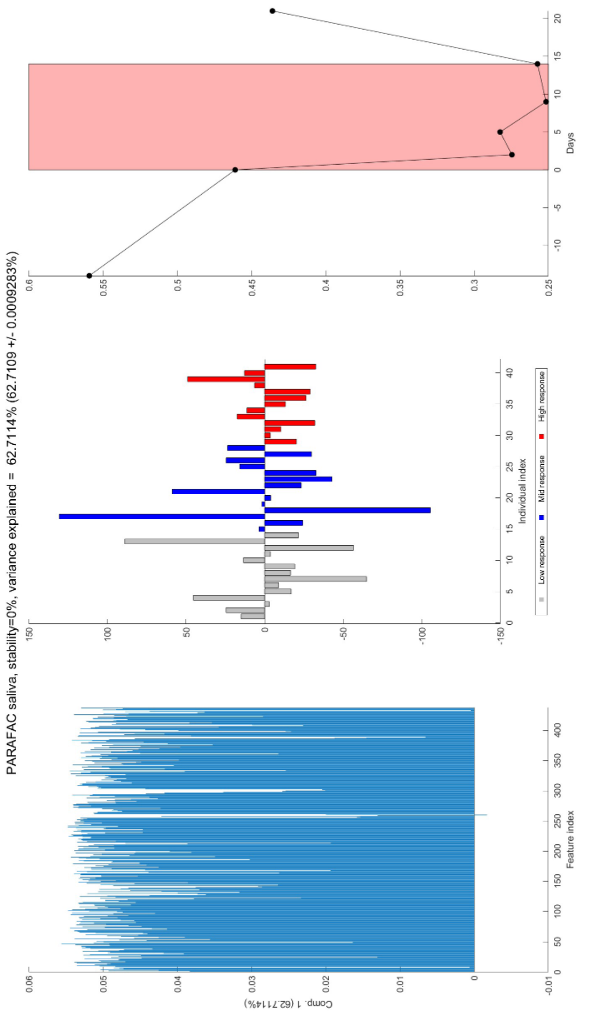


**Fig. A26** Overview of the PARAFAC model corresponding to the functional predictions of the samples obtained from the saliva microbiome. This model describes 62.7% of the variation in the data. In the left column, the ASV loadings are shown. In the middle column, the subject loadings are shown. In the right column the time loadings are shown. The subject loadings are coloured by their response group. In the time loading plot, the gingivitis intervention time points are shown with a red background.

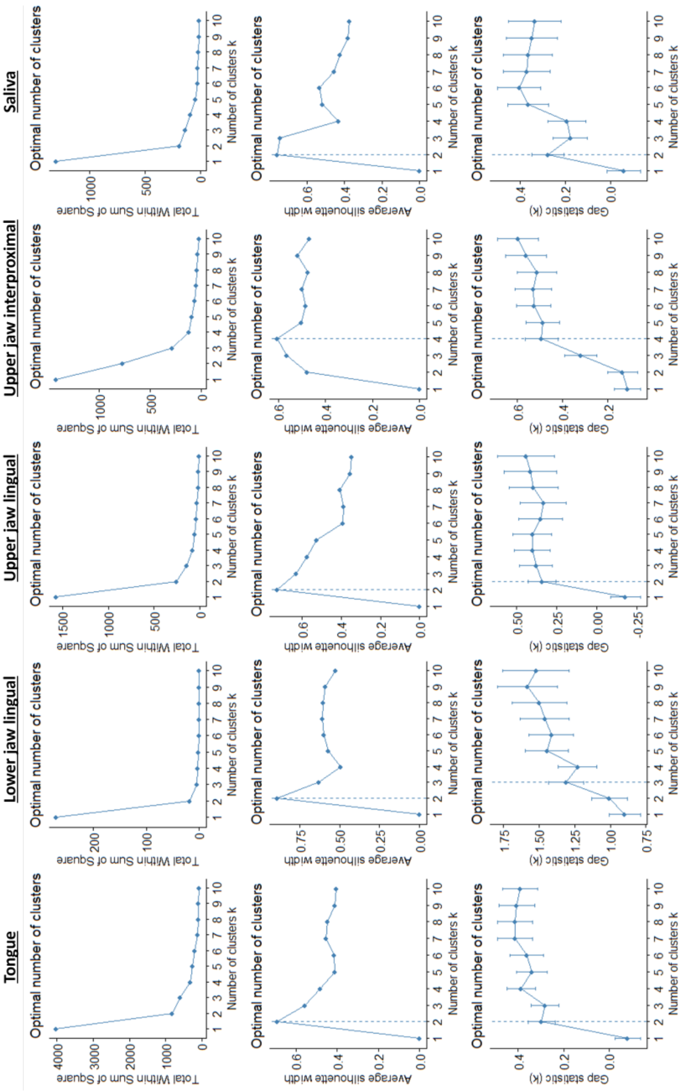


**Fig. A27** Overview of ASV clustering diagnostics per microbiome sample type. The top row shows the total within-cluster sum-of-squares metric. The middle row shows the average silhouette width metric. The bottom row shows the gap statistic. The optimal number of clusters is indicated with a dotted line and is generally followed if all metrics agree.

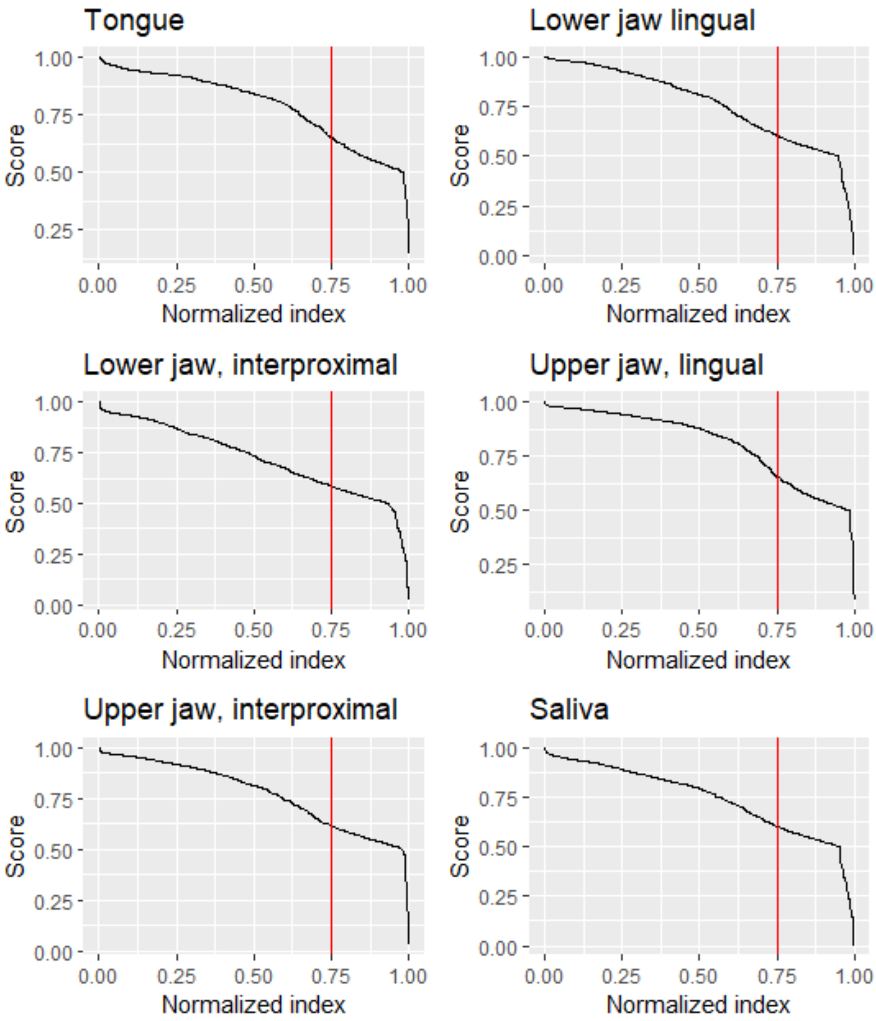


**Fig. A28** Overview of the integrated pathway element ranking for every functionally predicted microbiome sample type and the salivary metabolomics. Score is here defined as the average between the normalized variance explained and the normalized congruence loadings (see Methods). The ranking is cut of at 75% of its length, indicated by the red line, to obtain a meaningful separation between well-modelled pathway elements and averagely-modelled pathway elements.

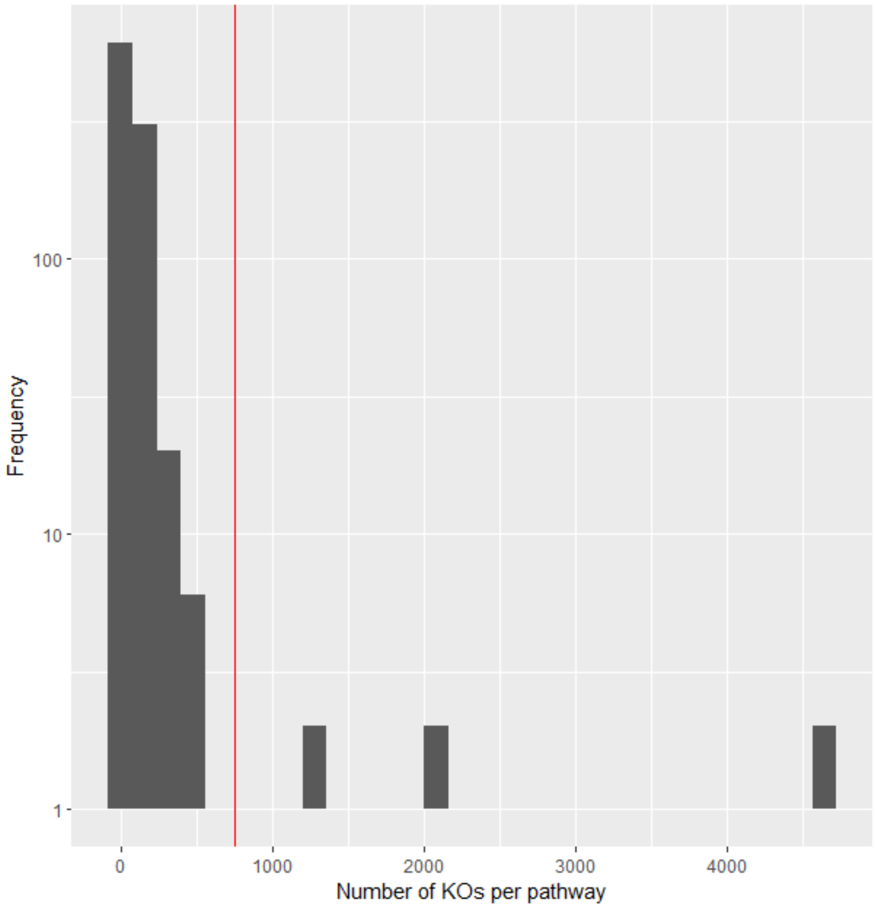


**Fig. A29** Histogram of KO terms per KEGG pathway. Note that one KO can map to multiple pathways. Pathways of more than 750 elements have been removed from the pathway enrichment analysis, indicated with the red line.

### Appendix B Supplementary Tables

**Table B1** Overview of the relationships between the subject loadings of the PARAFAC models and subject metadata. In the case of age, the Benjamini-Hochberg corrected p-values of the Pearson correlation test is shown. In the case of gender, the Benjamini-Hochberg corrected p-values of a Student's t-test is shown. Models are describing the microbiome data unless otherwise indicated. (*: $p \leq 0.05$; **: $p \leq 0.01$; ***: $p \leq 0.001$)

| Sample type | Component | Age | Gender |
|---|---|---|---|
| Tongue | 1 | 0.035* | 0.95 |
| Tongue | 2 | 0.94 | 0.90 |
| Lower jaw, lingual | 1 | 0.61 | 0.94 |
| Upper jaw, lingual | 1 | 0.85 | 0.70 |
| Upper jaw, lingual | 2 | 0.90 | 0.78 |
| Upper jaw, interproximal | 1 | 0.94 | 0.46 |
| Upper jaw, interproximal | 2 | 0.15 | 0.53 |
| Saliva | 1 | 0.15 | 0.68 |
| Saliva | 2 | 0.038* | 0.99 |
| Salivary metabolomics | 1 | 0.63 | 0.59 |
| Salivary metabolomics | 2 | 0.90 | 0.78 |

**Table B2** Overview of Benjamini-Hochberg corrected p-values of the permutation test for the differences between the mean sum of relative abundances between the low and high responders per ASV cluster per time point. This is based on 999 permutations of the response groups per time point/ASV cluster combination. (*: $p \leq 0.05$; **: $p \leq 0.01$; ***: $p \leq 0.001$)

| Sample type | ASV cluster | Visit 1 | Visit 2 | Visit 3 | Visit 4 | Visit 5 | Visit 6 | Visit 7 |
|---|---|---|---|---|---|---|---|---|
| Tongue | 1 | 0.045* | 0.10 | 0.18 | 0.30 | 0.37 | 0.29 | 0.33 |
| Tongue | 2 | 0.10 | 0.13 | 0.25 | 0.23 | 0.25 | 0.40 | 0.25 |
| Lower jaw, lingual | 1 | 0.020* | ≤0.001*** | ≤0.001*** | ≤0.001*** | ≤0.001*** | 0.062 | ≤0.001*** |
| Lower jaw, lingual | 2 | 0.020* | 0.19 | 0.42 | 0.41 | 0.55 | 0.35 | 0.19 |
| Upper jaw, lingual | 1 | 0.13 | 0.091 | 0.18 | 0.16 | ≤0.001*** | 0.011* | 0.029* |
| Upper jaw, lingual | 2 | 0.042* | 0.020* | 0.020* | 0.23 | 0.098 | 0.19 | 0.045* |
| Upper jaw, interproximal | 1 | 0.045* | 0.17 | 0.057 | 0.30 | 0.37 | 0.011* | ≤0.001*** |
| Upper jaw, interproximal | 2 | 0.039* | 0.13 | 0.47 | 0.036* | 0.020* | 0.13 | 0.12 |
| Upper jaw, interproximal | 3 | 0.39 | 0.25 | 0.47 | 0.20 | 0.091 | 0.10 | 0.35 |
| Upper jaw, interproximal | 4 | 0.16 | 0.15 | 0.10 | 0.18 | 0.020* | 0.20 | 0.29 |
| Saliva | 1 | 0.055 | 0.25 | 0.46 | 0.37 | 0.12 | 0.34 | 0.43 |
| Saliva | 2 | 0.029* | 0.15 | 0.25 | 0.15 | 0.30 | 0.28 | 0.42 |

**Table B3** Overview of Benjamini-Hochberg corrected Wilcoxon rank-sum tests between day 0 (the start of the gingivitis intervention) and day 14 (the end of the gingivitis intervention) for all identified ASV clusters per response group (see Table 2). (*: p≤0.05; **: p≤0.01; ***: p≤0.001)

| Sample type | ASV cluster | Low responders | Mid responders | High responders |
|---|---|---|---|---|
| Tongue | 1 | 0.21 | 0.88 | 0.25 |
| Tongue | 2 | 0.25 | 0.99 | 0.088 |
| Lower jaw, lingual | 1 | 4.0e-15*** | 4.0e-15*** | 7.2e-13*** |
| Lower jaw, lingual | 2 | 0.21 | 0.27 | 0.84 |
| Upper jaw, lingual | 1 | 0.70 | 0.069 | 0.002** |
| Upper jaw, lingual | 2 | 0.018* | 0.065 | 0.21 |
| Upper jaw, interproximal | 1 | 0.25 | 0.069 | 0.025* |
| Upper jaw, interproximal | 2 | 0.17 | 0.070 | 0.070 |
| Upper jaw, interproximal | 3 | 0.23 | 0.33 | 0.0039** |
| Upper jaw, interproximal | 4 | 2.6e-6*** | 1.3e-13*** | 1.3e-13*** |
| Saliva | 1 | 0.88 | 0.88 | 0.78 |
| Saliva | 2 | 0.21 | 0.56 | 0.12 |

**Table B4** Overview of the SetRank pathway enrichment result p-values per microbiome sample type, corrected for the multiple pathway membership of pathway elements. Pathways are reported here if they have at least one significantly ($p \leq 0.05$) enriched sample type, two well-modelled metabolites and two well-modelled microbiome molecular functions. (*: $p \leq 0.05$; **: $p \leq 0.01$; ***: $p \leq 0.001$)

| Pathway | Tongue | Lower jaw, lingual | Upper jaw, lingual | Upper jaw, inter-proximal | Saliva |
|---|---|---|---|---|---|
| Biosynthesis of amino acids | 0.19 | 0.027* | 0.072 | 0.47 | 0.30 |
| Biosynthesis of cofactors | 0.0045** | 0.18 | 0.012* | 0.089 | 0.016* |
| Biosynthesis of nucleotide sugars | 0.12 | 0.0012** | 0.0042** | 1.1e-4*** | 0.045* |
| Biotin metabolism | 0.059 | 0.37 | 0.17 | 0.054 | 0.045* |
| Carbon metabolism | 0.0052** | 0.25 | 1.0e-3*** | 0.11 | 0.010** |
| Glycerolipid metabolism | 0.55 | 0.49 | 0.037* | 0.71 | 0.60 |
| Inositol phosphate metabolism | 0.0037** | 0.39 | 0.58 | 0.17 | 0.28 |
| Lysine biosynthesis | 0.089 | 0.14 | 0.11 | 0.041* | 0.16 |
| Oxidative phosphorylation | 0.012* | 1.3e-4*** | 0.0041** | 7.9e-4*** | 4.1e-4*** |
| Purine metabolism | 0.039* | 0.020* | 0.035* | 0.0017** | 6.1e-4*** |
| Pyrimidine metabolism | 0.32 | 7.3e-4*** | 0.086 | 0.028* | 0.21 |
| Quorum sensing | 0.50 | 0.0019** | 0.031* | 0.028* | 0.0051** |
| Two-component system | 0.036* | 0.80 | 0.36 | 0.81 | 0.39 |

**Table B5** Overview of the number of variables in each dataset before and after variable selection. For the microbiome datasets: ASVs were removed if the sparsity was larger than 50% in all response groups. ASVs were also removed if they corresponded to chloroplast or mitochondrial sequences, as these were not relevant for the study. For the salivary metabolomics dataset: Metabolites were excluded if they contained more than 25% values below the detection limit. Additionally, xenobiotic compounds were manually assessed for occurence across response groups and selected if they were prevalent in most subjects. For the functionally predicted microbiome datasets: Predicted KEGG orthologous groups (KOs) were removed if they did not belong to pathways that the metabolites mapped to. KOs were also removed if the number of zeroes for all response groups was larger than 50%. Finally, KOs were removed if the sum-of-squares was lower than 0.025% of the total sum-of-squares in the dataset.

| Dataset | Sample type | #Variables after selection | #Variables before selection |
|---|---|---|---|
| Microbiome | Tongue | 106 | 2,892 |
| Microbiome | Lower jaw, lingual | 86 | 2,232 |
| Microbiome | Lower jaw, interproximal | 128 | 3,012 |
| Microbiome | Upper jaw, lingual | 65 | 2,253 |
| Microbiome | Upper jaw, interproximal | 93 | 3,719 |
| Microbiome | Saliva | 106 | 1,763 |
| Metabolome | Saliva | 400 | 499 |
| Functionally predicted microbiome | Tongue | 429 | 7,881 |
| Functionally predicted microbiome | Lower jaw, lingual | 465 | 7,995 |
| Functionally predicted microbiome | Lower jaw, interproximal | 439 | 8,094 |
| Functionally predicted microbiome | Upper jaw, lingual | 455 | 8,378 |
| Functionally predicted microbiome | Upper jaw, interproximal | 459 | 8,524 |
| Functionally predicted microbiome | Saliva | 437 | 8,017 |

**Table B6** Overview of sampling done per (anonymised) subject per sample type. Lowling: lower jaw, lingual. Lowinter: lower jaw, interproximal. Upling: upper jaw, lingual. Upinter: upper jaw, interproximal.

| ID | Tongue | Lowling | Lowinter | Upling | Upinter | Saliva | Metabolomics |
|---|---|---|---|---|---|---|---|
| 04KHQB | 7 | 6 | 5 | 7 | 7 | 7 | 5 |
| 07U72Y | 7 | 7 | 7 | 7 | 7 | 7 | 5 |
| 0Z20WA | 7 | 7 | 7 | 7 | 7 | 7 | 5 |
| 3830MH | 7 | 6 | 7 | 7 | 7 | 7 | 5 |
| 3CN8CB | 7 | 5 | 7 | 7 | 7 | 7 | 0 |
| 6U363N | 7 | 7 | 6 | 7 | 7 | 7 | 5 |
| 7OEMH5 | 7 | 7 | 7 | 7 | 7 | 7 | 5 |
| 7Y3TLP | 7 | 7 | 7 | 7 | 7 | 7 | 5 |
| 97Z53K | 7 | 7 | 7 | 7 | 7 | 7 | 5 |
| 9NCD7F | 7 | 7 | 7 | 7 | 7 | 7 | 5 |
| 9PSBPZ | 7 | 7 | 5 | 7 | 7 | 7 | 5 |
| BNXF8A | 7 | 7 | 7 | 7 | 7 | 7 | 5 |
| BOGUJ0 | 7 | 6 | 7 | 7 | 7 | 7 | 5 |
| COGRHY | 7 | 6 | 7 | 7 | 7 | 7 | 5 |
| D1NABQ | 7 | 7 | 7 | 7 | 7 | 7 | 5 |
| DDHX6K | 7 | 7 | 6 | 7 | 7 | 7 | 5 |
| DR0IOJ | 7 | 7 | 5 | 7 | 7 | 7 | 5 |
| EX26BI | 7 | 7 | 7 | 7 | 7 | 7 | 5 |
| GBLZO1 | 7 | 7 | 5 | 7 | 7 | 7 | 5 |
| GYIJCJ | 7 | 7 | 6 | 7 | 7 | 7 | 5 |
| IHNJDN | 7 | 7 | 7 | 7 | 6 | 7 | 5 |
| IICPTY | 7 | 7 | 7 | 7 | 7 | 7 | 5 |
| IOTBY5 | 7 | 7 | 6 | 7 | 7 | 7 | 5 |
| JN5XQ5 | 7 | 7 | 7 | 7 | 7 | 7 | 5 |
| KO9UPC | 7 | 6 | 7 | 7 | 7 | 7 | 5 |
| L5A913 | 7 | 7 | 7 | 7 | 7 | 7 | 5 |
| LZEI8O | 7 | 7 | 5 | 7 | 7 | 7 | 5 |
| MF64B7 | 7 | 7 | 7 | 7 | 7 | 7 | 5 |
| MZYB7A | 7 | 7 | 7 | 7 | 7 | 7 | 5 |
| O1HW8H | 7 | 7 | 7 | 7 | 7 | 7 | 5 |
| P4ZVS3 | 7 | 7 | 5 | 7 | 7 | 7 | 5 |
| RG524Q | 7 | 6 | 7 | 7 | 7 | 7 | 5 |
| RXOU2R | 7 | 7 | 7 | 7 | 7 | 7 | 5 |
| SMU8YW | 7 | 6 | 6 | 7 | 7 | 7 | 5 |
| TELB99 | 7 | 7 | 7 | 7 | 7 | 7 | 5 |
| W31R90 | 7 | 7 | 7 | 7 | 7 | 7 | 5 |
| WDSAGQ | 7 | 7 | 6 | 7 | 7 | 7 | 5 |
| XBBUT4 | 7 | 7 | 7 | 7 | 7 | 7 | 5 |
| YE3IMF | 7 | 7 | 6 | 7 | 7 | 7 | 5 |
| ZJ9VX5 | 7 | 7 | 7 | 7 | 7 | 7 | 5 |
| ZJWTJU | 7 | 7 | 6 | 7 | 7 | 7 | 5 |
| Total | 287 | 278 | 267 | 287 | 286 | 287 | 200 |

**Table B7** PARAFAC model statistics per dataset.

| Dataset | Sample type | Number of components | Variance explained (%) |
|---|---|---|---|
| Microbiome | Tongue | 2 | 31.5 |
| Microbiome | Lower jaw, lingual | 1 | 11.4 |
| Microbiome | Lower jaw, interproximal | 1 | 7.1 |
| Microbiome | Upper jaw, lingual | 2 | 19.7 |
| Microbiome | Upper jaw, interproximal | 2 | 16.5 |
| Microbiome | Saliva | 2 | 17.5 |
| Metabolome | Saliva | 2 | 22.6 |
| Functionally predicted microbiome | Tongue | 2 | 63.4 |
| Functionally predicted microbiome | Lower jaw, lingual | 1 | 49.6 |
| Functionally predicted microbiome | Lower jaw, interproximal | 2 | 72.4 |
| Functionally predicted microbiome | Upper jaw, lingual | 2 | 46.8 |
| Functionally predicted microbiome | Upper jaw, interproximal | 1 | 37.4 |
| Functionally predicted microbiome | Saliva | 1 | 62.7 |
